# Metagenome-based comparisons of decay rates and host-specificity of fecal microbial communities for improved microbial source tracking

**DOI:** 10.1101/2021.06.17.448865

**Authors:** Brittany Suttner, Blake G. Lindner, Minjae Kim, Roth E. Conrad, Luis M. Rodriguez, Luis H. Orellana, Eric R. Johnston, Janet K. Hatt, Kevin J. Zhu, Joe Brown, Konstantinos T. Konstantinidis

**Affiliations:** School of Civil and Environmental Engineering, Georgia Institute of Technology, Atlanta, GA 30332, USA; Ocean Science and Engineering, Georgia Institute of Technology, Atlanta, GA 30332,USA

**Keywords:** bioinformatics, comparative metagenomics, microbial ecology, water quality, public health, gut microbiome

## Abstract

Fecal material in the environment is a primary source of pathogens that cause waterborne diseases and affect over a billion people worldwide. Microbial source tracking (MST) assays based on single genes (e.g., 16S rRNA) do not always provide the resolution needed to attribute fecal contamination sources. In this work, we used dialysis bag mesocosms simulating a freshwater habitat that were spiked separately with cow, pig, or human feces to monitor the decay of host-specific fecal signals over time with metagenomics, traditional qPCR, and culture-based methods. Sequencing of the host fecal communities used as inocula recovered 79 non-redundant metagenome-assembled genomes (MAGs) whose abundance patterns showed that the majority of the fecal community signal was not detectable in the mesocosm metagenomes after four days. Several MAGs showed high host specificity, and thus are promising candidates for biomarkers for their respective host type. Traditional qPCR methods varied in their correlation with MAG decay kinetics. Notably, the human-specific *Bacteroides* assay, HF183/BFDRev, consistently under-estimated fecal pollution due to not being present in all hosts and/or primer mismatches. This work provides new insights on the persistence and decay kinetics of host-specific gut microbes in the environment and identifies several MAGs as putative biomarkers for improved MST.

**SYNOPSIS:** We track cow, pig, and human fecal pollution in lake water over time with metagenomics and benchmark these novel protocols against standard culture-based and qPCR tests for water quality monitoring.

## INTRODUCTION

Fecal indicator bacteria (FIB) are commonly used to assess microbial water quality and identify recent fecal pollution events. Because culture-based efforts to count FIB are ineffective for timely water management decisions, recent efforts have focused on rapid culture-independent qPCR methods targeting traditional FIB or new biomarkers (1–3). Members of the genus *Bacteroides*, or bacteriophages such as CrAssphage (4–7), are particularly suitable for microbial source tracking (MST) because they tend to co-evolve with the host, are among the most abundant genera in stool, have a narrow host range exclusive to warm-blooded mammals, and generally have poor survival rates outside their host (8). However, recent evidence also suggests the potential for *Bacteroides* to persist, and even grow under some environmental conditions (9, 10), which brings the assumptions about their survival outside of the host into question. Further, several studies report that these markers have some cross-reactivity with other (non-human) hosts (11–13) and may be too abundant in sewage for monitoring highly polluted waters (12). Clearly, more research is needed on the ecology of *Bacteroides* and other biomarkers (e.g., prevalence in human vs. animal hosts from different geographical regions), their persistence in the environment, and how well they correlate to risk of infection with enteric pathogens.

Further, the qPCR-based assays have their own (known) limitations (14). Specifically, it has been challenging to reliably compare the performance of different assays across different studies and environmental matrices because marker recovery efficiency and effect of PCR inhibitors can vary significantly among environmental samples (15, 16). Furthermore, even the most commonly used and studied human-associated markers (e.g., *Bacteroides* HF183) are not prevalent in all human populations worldwide (17, 18), which suggests no single qPCR marker is likely to be universally suitable for detecting human fecal contamination. Finally, fecal pollution of surface waters is often the result of a complex mixture of multiple inputs further complicated by environmental dispersion and deposition. The decay characteristics of different DNA markers is apparently of high importance for MST investigations and evaluation of the associated public health risk (19). Although an absolute gene count can be obtained via qPCR, estimates of the relative contribution of various fecal sources in the natural environment cannot be quantitative without this decay information. More comprehensive methods, such as metagenomics (20), can help to improve biomarker discovery and overcome several of the limitations described above.

Most research efforts utilizing metagenomics and next-generation sequencing (NGS) technologies thus far have focused on 16S rRNA gene amplicon sequencing to develop new biomarkers (21). However, the 16S rRNA gene is highly conserved across *Bacteria* and *Archaea*; as such, cross-reactivity with non-target hosts is common for all assays targeting even the most variable regions of the 16S rRNA gene (8, 22). Functional, protein-coding genes that are specific to a host’s unique gut physiology (e.g. host-microbe interactions) are likely more suitable targets for host-specific markers, but this represents a resolution level that 16S rRNA gene amplicon data cannot offer. Clearly, more research is needed to establish the best meta-omics and bioinformatic techniques as tools for identifying host-specific taxa and their genes for MST applications (23). Such studies will also establish whether or not metagenomic methods should be combined with conventional MST methods to obtain more accurate measures of fecal pollution in watersheds since qPCR provides absolute (vs. relative for typical metagenomics studies) abundances and generally has a lower limit of detection than metagenome shotgun sequencing (24, 25).

In this study, we used dialysis bag mesocosms simulating a fecal pollution event in a freshwater habitat and time-series metagenomics to track the decay of metagenome assembled genomes (MAGs) from human, cow, and pig fecal inputs over time. Additionally, we used traditional culture and qPCR-based MST markers and included a universal 16S rRNA gene qPCR assay for translating metagenome-based relative abundances to absolute abundances in order to directly compare against the traditional markers. Using the time-series abundance and cross-reactivity information, we identified ~12 MAGs as candidate MST biomarkers and compared their functional gene content to establish host-specific genomes and genes as potential targets for improved water quality monitoring assays.

## MATERIALS AND METHODS

### Lake water and fecal sample collection

Lake water samples were collected from Lake Lanier (Georgia, USA) in acid washed 10L carboys and transported immediately back to the lab and stored in the dark at 4 °C until mesocosm set up the following day (within 24 hours). Human fecal samples were collected from human volunteers who had not taken any antibiotics within the past one month before sample collection. Since human gut microbiomes are known to vary geographically, only samples from within the state of Georgia (USA) were used. All human subjects in the study provided informed consent and the study was approved by the Georgia Institute of Technology institutional review board (IRB) and carried out in accordance with the relevant guidelines and regulations. See supplementary methods for further details on sample processing.

### Mesocosm set-up

Sterile glass bottles were filled with 1.6 L of lake water and inoculated with feces to a final concentration of 2.5 g/L and shaken well to thoroughly mix the feces:lake water mixture before dispensing into dialysis bags. The dialysis bag’s pore size (6-8 kDa molecular weight cutoff) allows passage of small molecules and ions but prevents the passage of bacterial cells and viral particles. The dialysis bags were filled to a total volume of 110 mL (~21 cm length of 32 mm diameter dialysis tubing) and closed on both ends using polypropylene Spectra/Por clamps (Spectrum Laboratories). Enough dialysis bags were filled to sample each biological replicate in triplicate at each time point, i.e., 36 dialysis bags per host type (three technical replicates per three biological replicates at 4 sampling time points). Additionally, four uninoculated lake water negative control and two sterile milliQ water dialysis control bags were included for both of the two mesocosm experiment batches. The dialysis bags were suspended in ten-gallon aquarium tanks filled with lake water and stored in environmentally controlled rooms at 22 °C in the dark. A small water pump was included in each tank for aeration and nutrient distribution. A small headspace of air was left in each bag when sealing with the clamps so that they could float freely in the tanks.

### Mesocosm sampling

On the day of mesocosm set up, initial day zero (D0) reference community lake water samples were collected by filtering five separate 250 mL aliquots of uninoculated lake water onto 0.45 μm poly-carbonate (PC) membranes, three of which were stored at −80 °C in PowerFecal (Qiagen) 2 mL screw-cap bead tubes until ready for DNA extraction (within 1-3 months); two others were stored at −80 °C in sterile 2 mL screw-cap tubes filled with acid-washed 0.1 mm glass beads until ready for analysis following the EPA Method 1611 (26). Further, 100 mL of the lake water was filtered and cultured (in triplicate) on mEI medium following the EPA Method 1600 for culture-based enumeration of *Enterococci* (27). Finally, the feces:lake water mixtures were sampled following the same protocol for the un-inoculated lake water except using a 25 mL filter volume and 10-fold serial dilutions in 1X phosphate-buffered saline (PBS) for culture-based enumeration of *Enterococci* (27). All dilutions yielding measurements within the acceptable range of quantification were averaged to estimate CFUs/100mL of each biological replicate. To test for extraneous DNA and potential contamination from sample handling, 50 mL of sterile PBS was also filtered onto PC membranes and following the same DNA extraction at every sampling time point as described in the EPA method 1611 above to serve as a water sample filtration blank.

The qPCR markers used in this study are described in Table 1 and included the human-specific *Bacteroides* HF183/BFDRev (hereafter HF183; (2)), a ruminant-specific *Bacteroidetes* BacR (hereafter RumBac; (28)), human mitochondrial DNA (hereafter HUMmt; (29)), *Enterococcus faecalis* 16S rRNA gene (hereafter EF16S; (30)), the standard EPA Method 1611 assay targeting total *Enterococci* (hereafter EPA1611) and a universal 16S rRNA gene qPCR assay (hereafter GenBac16S; (31)) to normalize metagenome datasets for differences in microbial load. See supplementary methods for details on DNA extraction from feces and filters, conditions for qPCR assays and calculations for determining qPCR marker copy number and detection limits.

**Table 1:** qPCR markers used in this study and associated reference genomes. Host-specific MST markers include HF183, RumBac, and HUMmt; general FIB markers are EF16S and EPA1611. The GenBac16S assay was used for absolute quantification and LOD estimation for reference genomes in the metagenomes as described in the Materials and Methods section.

| Marker | Target | Reference | Reference genome | Accession |
| --- | --- | --- | --- | --- |
| HF183 | Human *Bacteroides* 16S | 2 | *Bacteroides dorei* CL03T12C01 | NZ_CP011531.1 |
| RumBac | Ruminant *Bacteroides* 16S | 48 | n/a | n/a |
| HUMmt | Human mtDNA NADH dehydrogenase subunit 5 | 49 | Human mitochondrion genome | J01415.2 |
| EF16S | *E. faecalis* 16S | 50 | *E. faecalis* ATCC29212 | CP008816.1 |
| EPA1611 | *Enterococcus* 23S | 46 | *E. faecalis* ATCC29212 | CP008816.1 |
| GenBac16S | Universal 16S rRNA | 51 | n/a | n/a |

### Metagenomic relative abundance estimation

Supplementary methods provide details on metagenome library creation and sequencing, detection of differentially abundant (DA) genes between host inocula or mesocosm time series samples, MAG recovery and dereplication at 95% average nucleotide level (ANI), and MAG annotation. To track relative abundance of MAGs or FIB reference genomes (Table 1) in dialysis bag mesocosm metagenomes, Magic-BLAST v1.4.0 ((32); options: -no_unaligned -splice F -outfmt tabular -parse_deflines T) was used to map metagenomic short reads to MAG contigs in order to express MAG abundance as average sequencing depth (base pairs recruited/genome length). Matches were filtered for single best alignments, using a minimum 90% query cover alignment length and 95% nucleotide identity of reads mapping against the reference genome (ANIr). In order to remove biases from highly conserved regions and contig edges, the 80% central truncated average of sequencing depth of all bases (TAD80) as described previously (33). MAG abundance in each metagenomic dataset (as % of total community) was calculated as the quotient of the MAG’s TAD80 value and the genome equivalents (GE) from MicrobeCensus (34).

### Approach for estimating metagenome limit of detection (LOD), absolute abundances, and decay rates

Reads belonging to the 16S rRNA gene were extracted with SortMeRNA v2.1 ((35); options: --log --fastx --blast 1 --num_alignments 1 -v -m 8336) and the SILVA 16S database dereplicated at 90% identity included with the program. The average 16S rRNA gene sequencing depth was estimated by summing the alignment length column from the SortMeRNA blast-like tabular output file and dividing by the average 16S rRNA gene length (1540bp). The average sequencing depth was then divided by the average genome sequencing depth from MicrobeCensus (34) to estimate the average 16S rRNA gene copy number per genome in the metagenome. The 16S rRNA gene copy number value was used to convert the GenBac16S qPCR count estimates to total cell density (number of prokaryotic cells per mL or mg) by dividing the qPCR count estimates by 16S rRNA gene copy number. With this information, it was then possible to estimate the theoretical LODs for a *Bacteroides* genome in each metagenomic dataset (in cells/mL) assuming at least 10% of a genome must be covered to reliably detect it in a metagenome (36) and that the average *Bacteroides* genome is 6.5 Mbp (34).

The absolute abundance (cells/mL) of fecal MAGs and reference genomes (Table 1) in the mesocosm metagenomes was estimated by multiplying its relative abundance (i.e. TAD80 >95% ANIr divided by GE) by the corresponding total cell density in each mesocosm metagenome. The same protocol was followed for the human mitochondrial reference genome for the HUMmt qPCR assay (Table 1), except sequencing depth was normalized using only the metagenome dataset size (in Gbp). Since there is no known reference genome for the RumBac assay, a 317bp contig from a cow fecal metagenome with a perfect match to the assay oligos (cow5_scaffold246842) was used as a proxy to estimate the absolute abundance as described above for genomes except no truncation was used when estimating sequencing depth (i.e. TAD100) and a 99% identity threshold for read mapping was used instead of 95% due to the high sequence conservation of the 16S rRNA gene relative to the rest of the genome (37, 38). The resulting sequencing depth value was divided by the 16S rRNA gene copy number for *Bacteroides* (n = 7) and genome equivalents (or GE) from MicrobeCensus (34) and subsequently multiplied by total cell density to estimate total number of *Bacteroides* cells per mL. The absolute abundances were used to calculate decay rates based on a first-order decay model, N_t_/N_0_ = 10^−kt^ (39). The time needed to produce a 2-log reduction in abundance (t_99_) was calculated using the decay constant (k) in the following equation, t_99_ = −2/k.

### Data Availability

Host fecal MAG assemblies and short reads for host fecal and mesocosm metagenomes have been deposited to NCBI databases under BioProject ID PRJNA691978, except the cow fecal metagenome short reads, which were deposited previously to the SRA database (BioProject ID PRJNA545149).

## RESULTS AND DISCUSSION

### Performance and decay of traditional culture-based and qPCR markers

Dialysis bag mesocosms simulating a natural freshwater environment were spiked with cow, pig, or human feces to represent a pollution event and monitored over time. Three biological replicate fecal samples were used per host and are referred to hereafter as hum1, hum2, hum3, cow7, cow8, cow9, pig7, pig8, and pig9 to indicate the specific individual host fecal sample that was used for DNA extraction and inoculation into the lake water mesocosms. H1, H2, H3, C7, C8, C9, P7, P8, and P9 hereafter refer to the feces:lake water mesocosm sample for each individual host (e.g. H1 refers to the lake water mesocosm spiked with feces from hum1). Mesocosm sampling occurred in triplicate on days 0, 1, 4, 7, and 14 (hereafter, D0, D1, D4, D7, and D14), which included qPCR analysis using the markers described in Table 1 and metagenome sequencing.

The qPCR markers were first tested against the host fecal DNA samples used as inocula to assess their sensitivity and specificity. The fecal DNAs were diluted 10-fold with water prior to running qPCR to reduce the effect of PCR inhibitors (see Supplementary Materials and Methods). All of the MST markers were not detected (ND) in any non-target hosts and none were quantifiable in the uninoculated lake water negative controls (Table S6). However, the human-specific HF183 marker was not detected in the hum2 fecal metagenome. The EPA Method 1600 culture-based test for *Enterococci* (EPA1600) showed that the dialysis bag mesocosms exceeded the EPA’s recreational water quality criteria (RWQC) of 36 CFU/100 mL throughout the entire duration of the cow and pig experiment and in all of the human timepoints except on D14 (Figure 1A). Furthermore, the EPA Method 1611 qPCR-based test for *Enterococci* (EPA1611) showed that all time-series samples that could be quantified exceeded the EPA RWQC of 103 calibrator cell equivalents (CCE) per 100 mL (Figure 1B). However, this assay was not detectable in the cow and pig samples by D14 and was only quantifiable in two of the three human samples on D1. Overall, the concentration of *Enterococcus* spp. was similar based on culture-based (EPA1600) and qPCR (EPA1611) assays and the first order decay rate constant was −0.20 d-1 for both methods (Table S5). Additionally, method blanks (sterile PBS filter controls) were included at each sampling point and analyzed according to the EPA1611 method and yielded no detectable amplification (data not shown), indicating no significant contamination during mesocosm sample handling.

**Figure 1:**
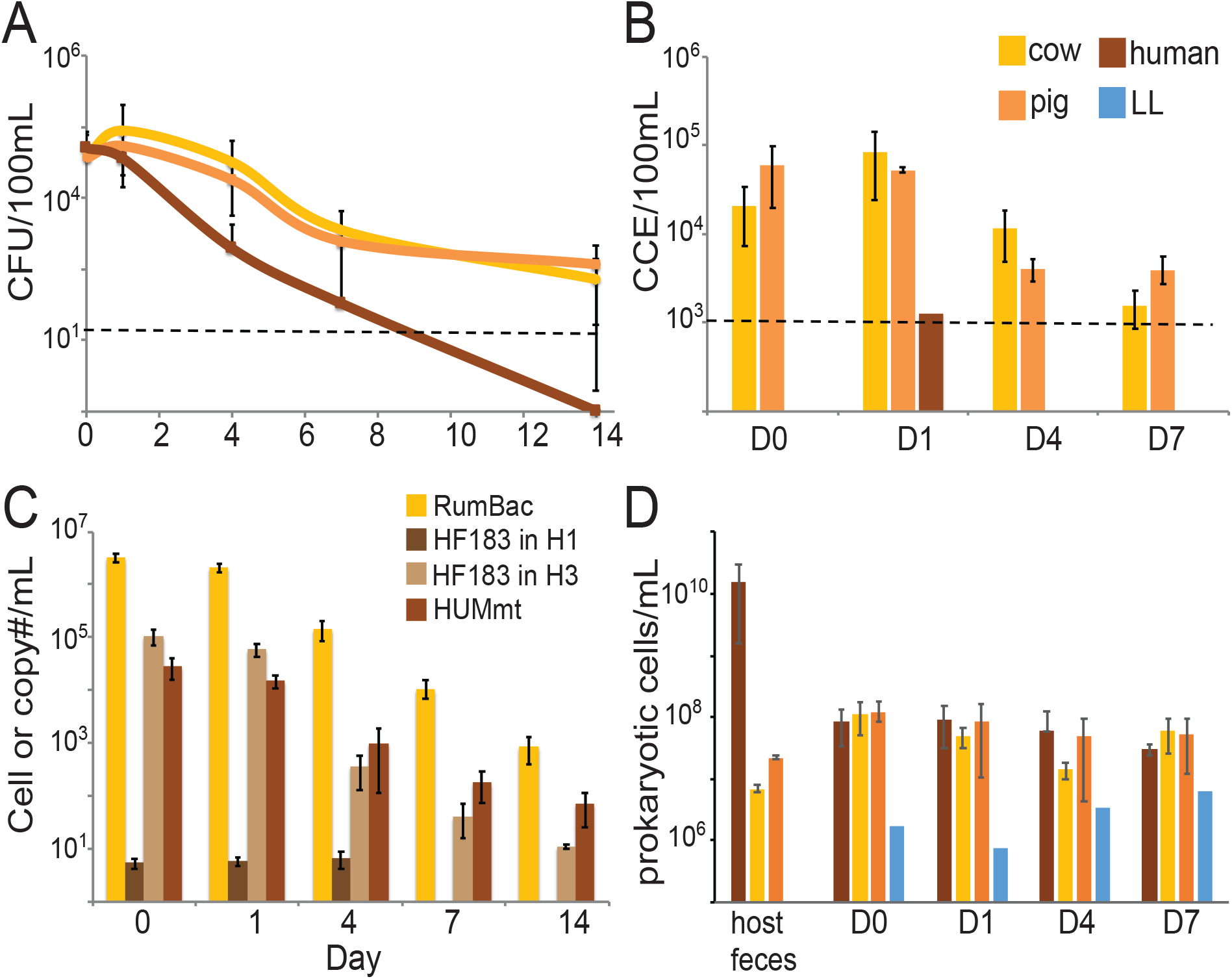
Traditional FIB, MST marker, and total bacterial cell abundances during the mesocosm incubations. (A) EPA Method 1600 culture-based enumeration of *Enterococcus*. (B) EPA Method 1611 qPCR-based enumeration of *Enterococcus*. Black dotted lines show the EPA’s recreational water quality criteria (RWQC) limit for impaired waters for each assay (CFU= colony forming units; CCE= calibrator cell equivalents). (C) Host-specific MST qPCR assays that could be detected in the dialysis bag mesocosms. The HUMmt is reported as #copies/mL and the rest are reported as #cells/mL. (D) Cell density in the mesocosms over time based on a universal 16S qPCR assay (GenBac16S). The average 16S rRNA gene copy number per genome was estimated from the corresponding metagenome for each sample by dividing average 16S gene sequencing depth by the average genome sequencing depth as described in the main text. In all figures, error bars are the standard deviation for averages that had more than three data points.

When tested in the time-series dialysis bag mesocosm samples, the qPCR gene copy estimates for all of the host-specific MST assays decreased with time and returned to very near or below the lowest concentration in the standard curves by D14 (~2.1 gene copies/μL DNA; Figure 1C). Consistent with the hum2 fecal DNA results, the HF183 marker was ND in any of the H2 mesocosm samples. The abundance of the HF183 marker in H3 mesocosms was two to four orders of magnitude larger than the abundances observed in H1 mesocosms (Figure 1C). Accordingly, only the H3 samples on D0 and D1 exceeded the quantitative microbial risk assessment (QMRA)-based water quality threshold of 41 copies/mL for HF183 as simulated for raw sewage in (40). Furthermore, the decay rates for the HF183 assay were 10-fold different in H1 and H3 (0.02 and −0.29 d^−1^, respectively; Table S5). The concentration of HF183 in H1 mesocosms was near the assay LOD (~5 cells/mL) at all time points, and thus there was no discernable decay for this marker resulting in the near-zero decay rate. The average gene copies/mL were consistent across the three biological replicates and were detectable until D14 for both the HUMmt and RumBac assays in human and cow mesocosms, respectively (Figure 1C).

The total cell density in the mesocosms based on the universal 16S qPCR assay (GenBac16S; Table 1) was ~10^8^ cells/mL at the start of all mesocosm incubations and tended to decrease with time, reaching ~1.5×10^7^ cells/mL by D7. The opposite trend was observed in the negative control bags, which started at ~10^6^ cells/mL and increased by nearly an order of magnitude by D7 (Figure 1D). These results indicated potential bottle effects during our incubations, which were assessed more fully by population genome binning of the D7 metagenomes as described in the Supplementary Results and Discussion. Notably, the bottle effect was consistent across all mesocosms and did thus, not affect the main results reported below.

### Taxonomic and phenotypic description of host-specific fecal MAGs

Host fecal reads were assembled into contigs with total length and N50 values ranging from 2.5×10^7^ to 1.3×10^8^ and 1,913 to 19,034 base pairs, respectively (Table S3; See also Supplemental Results for additional details on the metagenome datasets and community coverage). Contig binning from the inocula fecal datasets (not the mesocosm datasets) resulted in an initial set of 30 cow, 13 human, and 82 pig high quality MAGs. The MAGs were first dereplicated at 95% average nucleotide identity (ANI) within each host resulting in a new set of 18 cow, 13 human, and 50 pig MAGs. These MAGs were subsequently further de-replicated against the high quality MAGs from all three hosts and a collection of 477 Lake Lanier (LL) MAGs (33) to identify any MAGs that are non-host specific and/or found in the natural environment. This resulted in a final set of 17 cow, 13 human, and 49 pig high quality MAGs whose IDs are provided in Supplementary Data S1. MAGs were named according to the individual fecal sample from which they were originally assembled followed by the closest relative of the MAG and the lowest taxonomic rank the two share according to the MiGA TypeMat/NCBI database (p<0.1 threshold), i.e., C:class, O:order, F:family. G:Genus, S:Species. For instance, “cow4_20_Treponema_F” means MAG #20 assembled from cow4 fecal metagenome that had a *Treponema* sp. as the closest relative and was classified (at the lowest level with statistical confidence) to the family *Spirochaetaceae* (or, in other words, the MAG represents a novel genus and species of this family). Overall, the MAGs were highly host specific at the species level (ANI >95%) and there were only two instances (cow4_20_Prevotella_F and pig7_9_Tolumonas_C) in which a cow and pig MAG had ANI >95% with each other and were dereplicated into a single genomospecies (i.e., a cluster of genomes that is roughly equivalent to most named bacterial or archaeal species) and thus, were not used further as potential biomarkers. There was more overlap among hosts when evaluating the average amino acid identity values (%AAI; Figure S4) of their corresponding MAGs, revealing that these MAGs likely represent distinct but closely related species found in different hosts.

In all three host types, the majority of MAGs were classified at the class level as *Bacteroidia* (41%, 46%, and 33% for cows, humans, and pigs, respectively) followed by *Clostridia* (24%, 23%, and 31% for cow, humans, and pigs, respectively). In humans, the *Bacteroidia* MAGs were primarily assigned to the family *Bacteroidaceae,* whereas the cow MAGs were primarily from the *Prevotellaceae*. The majority of the pig MAGs could not be classified well below class level; i.e., they represented novel families (Supplementary Data S1). Consistent with their class level taxonomy, none of the host fecal MAGs (except pig6_25_Oscillibacter_O) were phenotyped as aerobes using Traitar (41) and the majority of MAGs were predicted to be anaerobes (100% human, 96% pigs, 82% cow; Figure S5). The oxidative stress enzyme catalase was not found in any of the cow or pig MAGs but was detected in two of the human MAGs (hum1_013_Akkermansia_G and hum2_003_Rubritalea_C). Glucose fermentation was the most common energy-yielding pathway in MAGs from all three host types (59% of cow, 71% of pig, and 100% of human MAGs). In addition to glucose fermentation, 44, 15, and 15 unique sugar substrates for growth were identified in the pig, cow, and human MAGs, respectively, with lactose being the most common in the pig and human MAGs (76% and 85% of total MAGs, respectively) and maltose being most common in the cow MAGs (82%). These results were also consistent with the DESeq2 analysis at the individual gene level (see Supplementary Results and Discussion).

### Decay kinetics of host fecal MAGs in the mesocosms

MAG abundance dynamics over the incubation time revealed that all 13 dereplicated human MAGs were detected in at least one human mesocosm, while only 13 out of 17 total cow and 41 out of 49 total pig MAGs were detected in all cow and pig mesocosms, respectively. For the conditions tested here, the majority of fecal MAGs from all three hosts were not detectable in the mesocosm metagenomes after D4 (Figures S7 & S8). Accordingly, it was only possible to estimate decay rates for 8 human, 3 cow, and 17 pig MAGs, respectively (Table S5) because at least three abundance data points (i.e., D0, D1, and D4) were required. For the MAGs with sufficient data points, the average log-2 reduction time (t_99_) was similar for cow and pig MAGs (4.5 and 5.6 days, respectively) but was higher for the human MAGs (average t_99_ of 14.3 days; Table S5). This result was largely consistent with a previous quantitative microbial risk assessment (QMRA) analysis that predicted that the gastrointestinal infection risk from sewage contamination in surface waters is not significant (<3% chance of infection) after 3.3 days (40) in accordance with the EPA risk threshold for bathing water (42). Specifically, Boehm et al, reported t_99_ of 1.4 d for protozoan, 2.5d for viral, and 4.7 d for *E. coli* O157:H7 and 11.8 d for *Salmonella*. Thus, the emerging 3-4 day rule of thumb for acceptable risk-levels seems to apply to many (but not necessarily all) fecal pathogens in aquatic environments. Importantly, the decay rate of the fecal MAGs is greater than or similar to those reported previously for several pathogens, suggesting that the MAGs may be suitable candidates for use as FIB with this respect. Consistent with this conclusion, none of the fecal MAGs were detected in any of the uninoculated lake water negative control metagenomes or matched closely any of the 477 LL MAGs, i.e., they are absent in the nearby natural ecosystem (Figure S11). Caution is needed, however, to extrapolate these results to all habitats, as some aquatic habitats or environmental conditions are known to support long-term survival of both pathogens and FIB (43). Furthermore, there are only a few studies to date reporting decay rates of viral markers and pathogens in the environment (44, 45) for evaluation against the MAG decay kinetics reported here; hence, viral pathogens may deviate from these decay patterns.

The human MAGs showed much higher individual host specificity than the cow and pig MAGs, i.e., MAGs assembled from an individual human fecal metagenome were always the most abundant in the mesocosms spiked with the feces from that individual and showed much lower abundances in the other two biological replicates (Figure S7A). In particular, among the hum2 MAGs, none were present in the H3 mesocosms and only two were detected in the H1 mesocosms; thus, none of the hum2 MAGs were selected as candidate biomarkers (see below). Therefore, targeting a single biomarker (whether it be a whole genome or qPCR assay) for MST can still be limiting due to the high individual variability observed in the human or animal gut, consistent with previous literature (46, 47), and the whole-community metagenomic approach employed here and/or targeting multiple biomarkers may be advantageous with this respect. Obviously, this limitation is not as important for MST in cases where the fecal input represents the composite excreta of many individuals such as in municipal sewage systems.

### Best-performing host fecal MAGs

Based on the decay and host specificity results, we identified five cow, three human, and six pig MAGs that were present in all three biological replicates of the same host type, were highly abundant on D0 (>0.1%) and were not detected in the metagenomes after D4 (Figures S7 and S8; Supplementary Data S1). We investigated these MAGs further as candidate biomarkers for MST. Notably, although most of the cow and pig MAGs are *Bacteroidia* and *Clostridia*, two of the cow biomarkers (cow4_001_Treponema_F and cow8_3_Treponema_F) were actually classified in the family *Spirochaetaceae*, while an *Actinobacteria* (pig4_16_Cellulomonas_C) and the archaeal phylum *Euryarchaeota* (pig4_38_Methanoplasma_F) were among the pig biomarker MAGs (Supplementary Data S1). These results suggest that biomarkers may be found in novel taxa not previously considered for MST.

Phenotype classification using Traitar (41) showed that none of the potential biomarker MAGs were aerobes or facultative anaerobes like the most commonly used FIB such as *E. faecalis* and *E. coli*, and all had primarily anaerobic phenotypes related to carbohydrate fermentation (Figure S5). Accordingly, the best gene targets for MST assay development at the individual gene level to detect relatively recent pollution events will likely be related to anaerobic functions specific to the different host types rather than the 16S rRNA gene, which has primarily been the target of most MST research to date. There were several functional genes that were significantly enriched in one host compared to the others and thus, could be targets for biomarker development (see also additional discussion in the Supplementary Material). These patterns, and the accompanying high host-specificity of the MAGs recovered, are presumably driven, at least to some extent, by the different selection pressures prevailing in the gut of each animal, as also indicated by the type of fermenters present in the different hosts.

#### Functional annotation for host-specific MAGs and gene functions

The 14 MAGs identified as potential markers based on their host sensitivity, abundance, and decay kinetics in the mesocosms showed no clear clustering based on the KEGG modules (48) found in their genome and no modules were clearly unique to a single host type (Figure S6). Thus, DESeq2 differential abundance (DA) analysis (49) based on reads mapped to assembled genes was used to identify specific functions that are enriched in the host fecal metagenomic assemblies. Of the 2,080 total KEGG functions identified, 177 were significantly DA with P_adj_ < 0.05 and log_2_ fold change (L2FC) > 3 using pairwise comparisons between human, cow, and pig fecal samples (Supplementary Data S2; Figure S13). Most of these gene functions were also recovered in the corresponding MAGs for the host type, further corroborating that the candidate host-specific MAGs are robust biomarkers (see Supplementary Results and Discussion for further details).

Most notably, seven genes for a type IV secretion system (T4SS) were highly abundant and specific to the cow gut metagenomes (Figure S13). Evidence has shown that T4SS proteins are important for shaping community composition in the gut (50), which suggests these proteins could be viable targets as host-specific markers. These results were also consistent with another study using competitive DNA hybridization to survey metagenomes and found host-specific sequences related to secretion and surface-associated proteins (51). Furthermore, some of the DA KEGG functions offered new insight on the fermentation pathways that distinguish cows and pigs. Fumarate reductase subunit D (*frdD*), which is associated with the primitive electron transport chain (ETC) of some fermenters (52), was more abundant in cows. The pig samples were instead enriched for two genes associated with butyrate-producing fermentation (*atoA*; acetoacetate CoA-transferase beta subunit, *bcd*; butyryl-CoA dehydrogenase) as well as the gene associated with H_2_-producing fermentation (*porD*; pyruvate ferredoxin oxidoreductase delta subunit) (Supplementary Data S2). These results indicated that fermenting microbes inhabiting the cow and pig gut carry out different strategies to sink excess reducing equivalents (the primitive ETC or H_2_, respectively). However, these trends were not discernable for the human samples as fewer genes overall tended to be significantly enriched in human inocula, which could be the result of sampling limitation (only 3 human fecal samples were compared against 6 cow and 6 pig samples) and the higher inter-person diversity described above.

### Decay of potential biomarker MAGs vs. reference FIB genomes in the mesocosms

The absolute abundances (cells or viral particles/mL) of common FIB and MST biomarkers over time were also compared to the candidate host-specific biomarker MAGs. The former biomarkers included reference genomes associated with the qPCR assays used in this study (Table 1) as well as genomes of the common commensal *E. coli* HS (NC_009800.1) and CrAssphage (JQ995537). *E. faecalis* and *E. coli*, despite being “gold standard” FIB, performed worse than the MST markers described here. *E. faecalis* was not detected in any human feces or mesocosm samples by qPCR or metagenome-based methods and its abundance was too low in the cow and pig feces to be detected upon dilution in the lake water mesocosms (data not shown). Hence, this organism would not be able to indicate fecal contamination for any of the hosts in this study. *E. coli* was detected in all of the host mesocosms and persisted for ~1 week (Figure S16A), longer than the presumed fecal contamination risk of 4 days described above, and maintained higher abundances over time compared to the fecal MAGs (Figure 2). Although it is well known that *E. coli* and *E. faecalis* are not host-specific, and thus their usefulness for MST may be limited, these results confirmed our expectations and provided further evidence against the use of these organisms as FIB and highlight the need for improved standard indicators.

**Figure 2:**
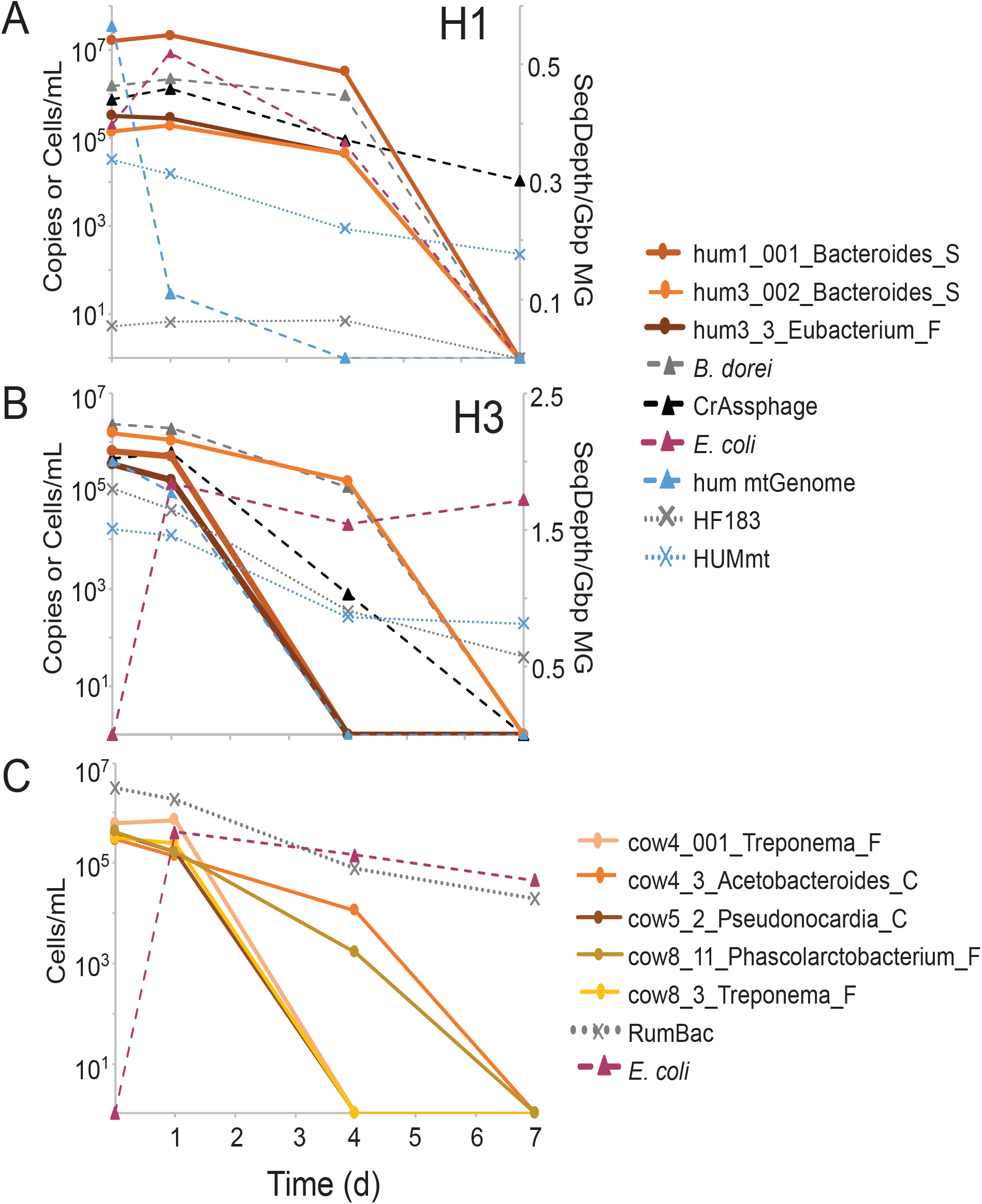
Compare absolute abundances of putative biomarker MAGs, traditional FIB and MST qPCR markers in (A) H1 mesocosms, (B) H3 mesocosms, and (C) the average of all 3 biological replicates of the cow fecal mesocosms. Absolute abundances (gene copies, cells or viral particles per mL) were determined for all targets except for the human mitochondrial genome (hum mtGenome), which is expressed as relative abundance (sequencing depth per Gbp metagenome) and is shown on the secondary axis for (A) and (B). MAGs are represented by solid lines with circle markers. Reference genomes are represented by dashed lines with triangle markers and include *Bacteroides dorei*, CrAssphage, *E. coli*, and the human mtGenome. The qPCR assays are represented by dotted lines with X markers and included the human-specific and ruminant specific *Bacteroides* assays (HF183 and RumBac, respectively) and the human mtDNA assay (HUMmt; reported as copies/mL). The human mesocosms are plotted separately because they were more variable among each other compared to the cows and also because neither *B. dorei*, CrAssphage, or HF183 were detected in any of the H2 mesocosms. Thus, H2 is not shown here.

The *B. dorei* and CrAssphage genomes were not detected in any cow or pig mesocosm metagenomes and they also had similar decay profiles to the human fecal MAGs (Figure 2A & B); except the CrAssphage genome abundance increased from D0 to D1, whereas the *B. dorei* genome (and fecal MAGs) abundance consistently decreased with time, which could possibly indicate a predatory relationship between these two microbes (CrAssphage is predicted to be a *Bacteroides* bacteriophage). Further, consistent with the qPCR results, neither of these genomes were detected in any of the H2 mesocosm metagenomes. *Bacteroides* abundance based on the HF183 qPCR assay tended to be lower than the MAGs and *B. dorei* reference genome (see below). The human mitochondrial genome (mtGenome) was detected in all three human mesocosm metagenomes until D7 and showed a steady decay in abundance with time (Figure 2A & B and S16B), consistent with the HUMmt assay, which was detectable by qPCR until D14 (Figure 1C). The cow fecal MAGs were all ND by D7 and decayed faster compared to the *E. coli* reference genome and *Bacteroides* abundance based on the RumBac qPCR assay (Figures 2C, S5 and S6).

#### Correlation of MST qPCR markers to their metagenome counterpart

In order to more precisely evaluate the performance of the metagenome-based results against those of traditional qPCR assays, absolute abundances (expressed as cells/mL) of the RumBac and HF183 *Bacteroidetes* assays were compared to the abundance of the corresponding reference genome in the mesocosm metagenomes. The correlation between *Bacteroides* abundances based on qPCR and metagenomes estimates was not consistent between the two assays (Figure 3A & B; R^2^=0.18 and 0.76 for HF183 and RumBac, respectively). The RumBac qPCR assay tended to give higher abundance estimates (linear regression slope = 0.16) than its metagenome counterpart (Figure 3B). The HF183 qPCR assay consistently gave lower estimates of *Bacteroides* abundance in the human mesocosms (linear regression slope= 10.26), especially in H1, in which the HF183 qPCR assay estimated only about 6 *Bacteroides* cells/mL in the mesocosms on D0, D1, and D4, well below the theoretical LOD for *B. dorei* in the metagenomes (~3×10^4^ cells/mL; see Materials and Methods for LOD estimation). However, the *B. dorei* reference genome was well above this concentration based on metagenome abundance (Figure 3A). Further investigation showed that this was presumably caused by mismatches of the forward HF183 primer to the dominant *Bacteroides* strains present in the host fecal inocula (Figure S15). Specifically, the short reads from the fecal inoculum were searched against the 16S rRNA gene of the reference *B. dorei* strain (which contains a perfect match to the HF183 assay primers and probe) to calculate its 99% identity truncated average sequencing depth (TAD80). For both hum1 and hum3 fecal metagenomes (there was no detection in hum2), the sequencing depths of the probe and reverse primer were similar to the overall average sequencing depth for the entire 16S rRNA gene (at about 42.0 and 247.0 for hum1 and hum3, respectively). However, the sequencing depth of the forward primer region was 0 in hum1 and ~40 (6x less than the average) in hum3. Furthermore, we manually checked the metagenomic reads for perfect matches to the HF183 forward primer and found none in hum1 and only 17 in hum3, suggesting that this region is not present in the dominant *Bacteroides* strain that was assembled from each host.

**Figure 3:**
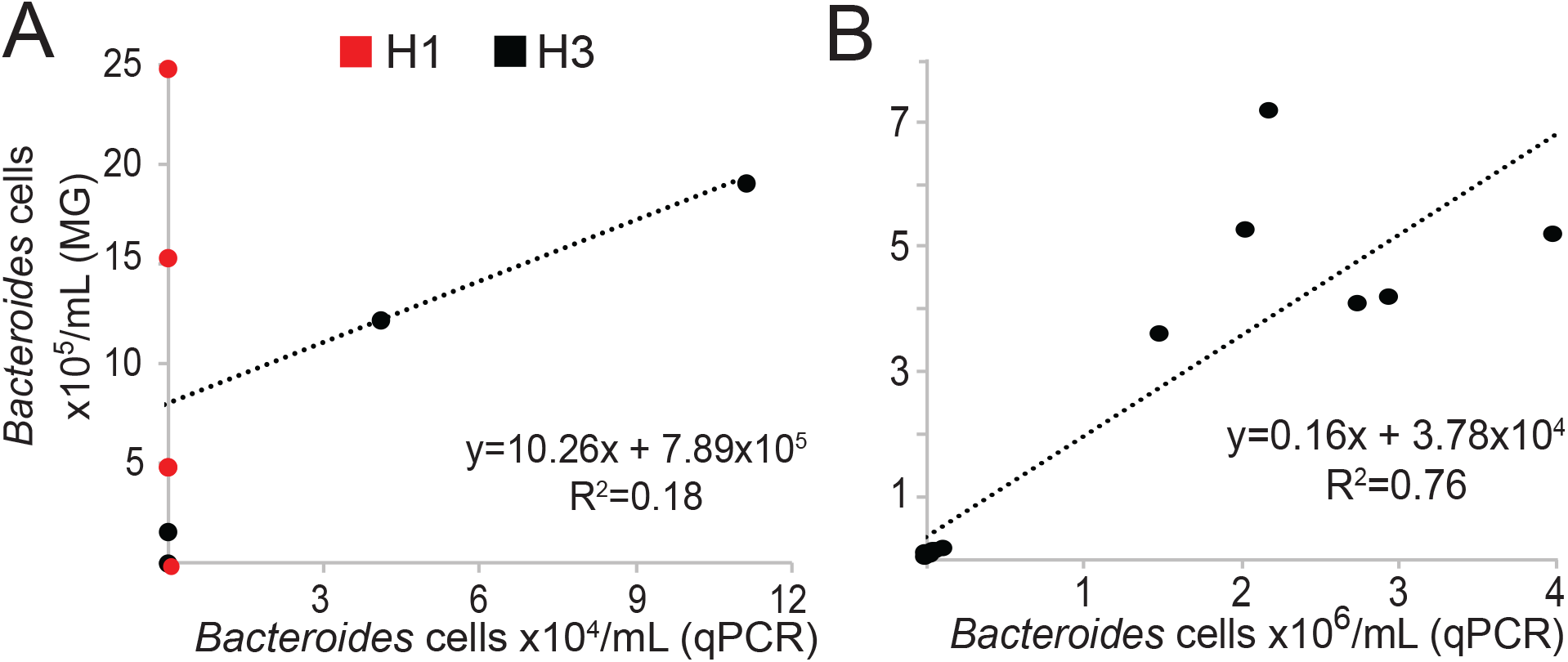
Correlation between qPCR and metagenome-based abundance estimates of MST markers and their reference genome counterparts. (A) Human-specific *Bacteroides* 16S (HF183) versus the absolute abundance of the reference genome *B. dorei* in the human mesocosm metagenomes. (B) Ruminant-specific *Bacteroides* 16S (RumBac) versus the absolute abundance of a contig recovered from the cow fecal inocula metagenomes carrying a perfect match to the RumBac assay in the cow mesocosm metagenomes. Absolute abundances for (A) and (B) are expressed as the number of *Bacteroides* cells/mL.

Thus, our evaluation of traditional qPCR assays revealed several of the known limitations of this approach such as mismatches of the PCR primers against the taxa present in the sample (Figure S15), and lower decay rates of short DNA fragments (~120bp) targeted by PCR relative to whole cells or the whole chromosome (see also Supplementary Results and Discussion). It is important to note, however, that the PCR primer limitation is not expected to be as pronounced in cases where the fecal input represents many individuals due to the high inter-individual variability in the microbiome. In such cases, the PCR markers such as the HF183 are expected to perform well for their purposes, as previously noted (40). Furthermore, there was some overlap among the different host fecal MAGs at the genus level (>65% AAI; Figure S4), which, most likely, accounts for the cross-reactivity commonly observed for the various 16S qPCR assays targeting *Bacteroidales* at above the species level. (8, 16, 22).

### Using metagenomic methods for microbial source tracking

Collectively, our findings suggest that the use of metagenomic methods to identify host-specific MAGs and detect and track these MAGs in an environmental-like system is highly promising and circumvents several of the limitations of traditional methods. Considering the high individual host variability, especially among human hosts, more work is needed to characterize the geographic stability of the putative biomarkers of human or animal hosts reported here and the degree of their biogeography. All of the cow and pig fecal samples used here were from the same farm in northern Georgia (USA) for the convenience in obtaining these inocula as well as technical limitations in running the mesocosm incubations with a larger number of samples. Thus, it will be important to determine if these MAGs are present in animals from other herds across broader geographical regions. Many recent studies have made considerable effort to sequence metagenomes and/or assemble MAGs from cow rumen (53–55) as well a pig (56, 57) and chicken guts (58). However, this information has not yet been synthesized together for MST marker development. Future work should leverage these datasets to improve comparative functional gene analysis along with decay information to search for better DNA markers. Furthermore, as high-throughput sequencing becomes more affordable and routine, it may be possible to directly assess MST markers (and even pathogens) in environmental metagenomes. To make regulatory standards based on metagenome data, calculating absolute abundances of indicators (or pathogens) will be necessary. The methodologies proposed here should be helpful in these directions.

## Acknowledgments

The authors would like to thank Dr. Robert Dove from the University of Georgia-Athens for providing the cow and pig fecal samples. This work was supported by the US National Science Foundation, award numbers 1511825 (to J.B and K.T.K) and 1831582 (K.T.K.), and the US National Science Foundation Graduate Research Fellowship under grant number DGE-1650044 (to B.S). The funding agencies had no role in the study design, data collection and analysis, decision to publish, or preparation of the manuscript.

