## Supplementary Information for "Metagenome-based comparisons of decay rates and host-specificity of fecal microbial communities for improved microbial source tracking"

**Supplementary Materials and Methods**

*Fecal sample processing:*

Remel fecal collection kits (ThermoFisher) were provided to human volunteers, who were instructed to store their sample at 4 °C and return within two days after fecal collection. Cow and pig fecal samples were collected within six hours of defecation from animals at the University of Georgia Athens Department of Animal and Dairy Sciences farms. A portion of each freshly-excreted fecal sample was preserved for DNA extraction by adding 1:1 volume feces to one mL sterile distilled water for a total of two mL and stored at 4 °C until processed in the lab. Five mL of lysis buffer (Qiagen PowerBead solution) was added to the water:feces mixture, vortexed for 30 seconds, then spun for three minutes at 1500rpm to create a homogenized fecal slurry. One to two mL of the slurry was pipetted into two mL cryo-vials and stored at -80 °C until ready for DNA extraction. A separate portion of each freshly-excreted fecal sample was also persevered for inoculation into mesocosms by adding 1:1 volume feces to 15 mL sterile Cary-Blair media for a total of 30 mL and stored at 4 °C until mesocosm set-up (within two days of sample collection).

*DNA extraction from feces and filters:*

DNA was extracted from homogenized fecal slurries for the quantification of MST qPCR markers and metagenome sequencing. The Qiagen PowerSoil kit was used for the cow, pig, and human fecal slurries following a modified Human Microbiome Project protocol for stool samples described previously (1). Two separate DNA extraction protocols were used for the PC filters (i.e., two filters per dialysis bag sample were used for DNA extraction with two different methods). One PC filter was used for quantification of MST qPCR markers and metagenome sequencing and DNA was extracted using the Qiagen PowerFecal kit following the manufacturer’s instructions except mechanical cell lysis was performed by bead beating in two 1-minute intervals using the Biospec Mini-Beadbeater-24. The other PC filter was used for DNA extraction and enumeration of total *Enterococci* following the EPA Method 1611 (2). This method was designed for rapid and simple water quality monitoring and does not include any chemical precipitation or clean-up steps. Therefore, this method results in a more “crude” DNA extraction that is not suitable for most qPCR and metagenomic methods.

*qPCR for common MST markers:*

All qPCR reactions were run using an Applied Biosystems 7500Fast thermocycler and the cycling parameters were as follows: 2 min at 50 °C, 10 min at 95 °C, and 40 cycles of 15 sec at 95 °C and 60 sec at 60 °C. The EPA Method 1611 assay was run using the standard calibrator cells and the ΔΔCt quantification method following the protocol described in (2)**.** All other qPCR assay reactions used two ul of extracted DNA as template in 20 μL qPCR reactions with the TaqMan Universal PCR Master Mix (Applied Biosystems). Template DNAs were run undiluted or diluted 10-fold (i.e. to remove the effect of PCR inhibitors) depending on the expected marker concentration and quality of each sample. Initial tests of running 10-fold diluted and undiluted fecal DNAs showed that Ct values differed by less than 3x and indicated the presence of PCR inhibitors. Thus, only the results from the diluted samples were used in downstream analyses. No significant PCR inhibitors were detected in tests with the mesocosm DNAs; thus, these were run undiluted. The Taqman (i.e., 5’ hydrolysis) probe and primer concentrations for each assay are listed in Table S4 and only the RumBac assay reactions included 10 μg bovine serum albumin (BSA). All samples were run in triplicate on 96-well plates and each run included triplicate no template controls (NTC). In all plates, the NTCs had no detectable amplification or were at least 1 Ct value larger than the average Ct value of the lowest concentration in the standard curve. The amplification threshold for the GenBac16S, EF16S, and HUMmt assays was set to 0.2 ∆Rn units and to 0.03∆Rn units for the HF183 and RumBac assays.

Standard plasmids were used for absolute quantification. Target sequences were ligated into the pCR™2.1-TOPO® TA vector and cloned using One Shot® Chemically Competent *Escherichia coli* TOP10 and the TOPO®-TA cloning kit (Invitrogen), following manufacturer’s instructions. The qPCR amplicon sequences were used as the target sequence in the standard plasmids for the HF183, RumBac, and HUMmt assays, while a full-length *E. faecalis* 16S rRNA gene was used in the standard plasmid for the EF16S and GenBac16S assays. Genomic DNA from *Bacteroides* sp., Strain 1_1_6, HM-23D (obtained through BEI Resources, NIAID, NIH as part of the Human Microbiome Project) was used at the template DNA for generating the HF183 qPCR amplicon for ligation with standard plasmid. Standard plasmids were isolated using the QIAprep Spin Miniprep Kit (Qiagen) following the manufacturer’s instructions and quantified using the Qubit HS DNA kit. Seven, 10-fold serial dilutions (10^6^ to 10^0^ copies per reaction) of qPCR standard plasmids were run in triplicate on every 96-well plate. Marker concentrations were determined using the corresponding calibration curve from each plate because no more than four plates were run per assay. Details on the standard curves and average assay efficiencies for each assay are provided in Table S4.

*Determination of qPCR marker copy number and limit of detection:*

Marker copy number per qPCR reaction was calculated for all samples using the linear fit of log-transformed standard copy number versus threshold cycle (Ct). A marker was considered not detected (ND) for any sample that did not return a Ct value in two of the three triplicates and was considered detectable but not quantifiable (DNQ) if it returned an average Ct value that was above the average Ct value of the lowest concentration in the standard curve (i.e., the limit of quantification), which was ~2.1 counts/μL DNA for the HF183, RumBac, EF16S, and HUMmt assays and 168 counts/μL DNA for the GenBac16S assay. The copy number per uL of each standard plasmid was calculated based on the concentration (in ng/uL) and assuming 660 g/mole bp DNA. The number of gene copies in each detectable sample was averaged, normalized by the volume of DNA per reaction, multiplied by the DNA elution volume (100 μL) and then divided by the total water filter volume (mL) or the mass of feces used in the initial DNA extraction (mg). The gene count estimates for the HF183 and RumBac assays were converted to absolute abundances assuming 7 copies of the 16S rRNA gene per genome. Because these assays are expected to target several related species of the *Bacteroides* genus (HF183;(3)) and *Bacteroidetes* phylum (RumBac;(4)), their abundances are expressed roughly at the genus level (i.e., as the number of *Bacteroides* cells/mL) for both the qPCR and metagenome abundances (described below).

*Metagenome library sequencing, quality assessment and analyses:*

Metagenome sequencing libraries were prepared using the Illumina Nextera XT kit and sequenced on the HiSEQ 2500 instrument as described previously (5). Three additional negative control libraries that were not included in the initial HiSEQ 2500 run were sequenced later on the NovaSEQ 6000 S4 platform (i.e. animal_LL_D1, D4, and D7) The two D0 negative control libraries (i.e. human_LL_D0 and animal_LL_D0) were also re-sequenced on the NovaSEQ for quality control comparisons to HiSEQ data. Short reads were passed through quality filtering, trimming and assembly using MiGA (6) with the default settings (PHRED score cutoff of 20, only retain read pairs with both sisters ≥ 50bp after trimming, and assembly with IDBA-UD (7) using kmer values ranging from 20 to 80). MASH v1.0.2 (options: -s 100000;(8)) was used to determine whole-community similarity between metagenomes in a reference-independent approach. The MASH distances were used for ANOSIM, ADONIS (compared by host type and sampling day), and non-metric multidimensional scaling (NMDS; number of dimensions =4) with the metaMDS function in the R package vegan v2.5-6. Average community coverage and diversity were estimated using Nonpareil v3.0 (9) with kmer kernel and default parameters. Average genome size and genome sequencing depth (i.e. average sequencing depth of single copy genes) were estimated in each metagenomic sample using MicrobeCensus v1.0.6 with default parameters (10).

*Gene functional annotation and determination of differentially abundant (DA) gene functions in host fecal and D7 metagenome assemblies:*

Open reading frame (ORF) prediction from assembled contigs was performed using Prodigal (Hyatt et al. 2010) as implemented in MiGA (6). Resulting amino acid sequences were searched against the KEGG ortholog profile HMM database (KOfams) using KoFamScan v1.2.0 (11) with the ‘prokaryote’ database and using the parameter ‘-f mapper’ to provide only the most confident annotations (i.e., ORFs assigned an individual KO). Orthologies were matched to their corresponding functional annotations using a parsed version of the KEGG orthology table (‘ko00001.keg’; https://github.com/edgraham/GhostKoalaParser). Sequence coverage of each gene was determined by mapping metagenomic short reads against the corresponding ORFs for each sample using Magic-BLAST v1.4.0 (12). The Magic-BLAST outputs were filtered for best match*,* 90% query cover alignment length, and a minimum read length of 50 bp. These read counts were used to determine DA functional annotations in samples grouped by host type (i.e. pairwise comparisons of human, cow, and pig fecal samples) and all host fecal metagenomic samples vs. D7 mesocosm metagenomes using the negative binomial test and false discovery rate (P_adj_ <0.05) as implemented in DESeq2 v1.4.5 (13). DA functional annotations with Log_2_ fold change (L2FC)>|3| for the host only and L2FC>|6| for the host vs. D7 comparisons were summarized into several hierarchical ranks including metabolic pathways and individual protein families based on the KEGG classification system (Supplementary Data S2 and S3). A larger L2FC cutoff was used for host fecal vs. D7 comparison so that the number of DA functions retained were feasible for manual inspection (i.e. <600 functions). Read counts for each summarized functional category were converted to genome equivalents (GE) by dividing by the average genome sequencing depth as determined using MicrobeCensus v1.0.6 (10). Each category was divided by the average GE across all samples to provide unbiased counts for visualization purposes.

*Binning of fecal and D7 metagenomes and dereplication of MAGs:*

The host fecal and D7 dialysis bag metagenome assemblies were used for population genome binning with MaxBin 2.2.4 (14) and MetaBat 2.12.1 (15) with default settings. Only contigs longer than one kbp were used for binning. Fecal MAGs from the two algorithms within each host type were combined and de-replicated with DAS Tool 1.1.0 (16), and only resulting MAGs with contamination <5% or MiGA quality score >50 were retained for further analysis. This collection of high-quality host fecal MAGs was also dereplicated against each other (i.e., across each host type), along with a collection of 477 Lake Lanier (LL) MAGs (17), and the non-aggregated set of D7 MAGs (i.e. the total set of MAGs resulting from both MetaBat and MaxBin) using the MiGA derep workflow (6). That is, in cases where two MAGs shared >95%ANI, only the higher quality MAG was retained for further analysis. The average amino acid identities (AAI) calculated by MiGA were used to generate heatmaps with the seaborn library in python3.

*Taxonomic and functional annotation of MAGs:*

Taxonomy was assigned to the MAGs using the MiGA assign_taxonomy workflow and the TypeMAT/NCBI database. MAG phenotypes (aerobe, anaerobe, fermentation type) were assigned using Traitar v1.0.4 and the prediction results from the phypat classifier model only (18). Functional annotations were assigned to genes of MAGs identified as potential biomarkers (see following section) using KoFamScan v1.2.0 (11) and the Swiss-Prot (19) database as implemented in MicrobeAnnotator (Ruiz-P et al. *in review*; https://github.com/cruizperez/MicrobeAnnotator) with the -m sword and --light options.

**Supplementary Results and Discussion**

### *General description of metagenome samples and community coverage:*

### A total of 56 metagenomes, ranging in size from 3.2 to 35.4 million reads (0.4 to 3.8 Gbp) were determined from 36 dialysis bag mesocosms (feces:lake water mixture), 5 lake water negative controls, and 15 host fecal samples (Table S2). Fecal metagenomes from 3 cow and 3 pig individuals were also included in this study (in addition to the 3 cow and 3 pig fecal samples used as inocula in the mesocosms) because considerably less fecal metagenomes have been sequenced for these hosts compared to humans and less is known about their gut microbiome diversity. The average fraction of the total community covered by our sequencing efforts as determined by Nonpareil (9) was consistent across the biological replicates for the fecal metagenomes (Table S1). Given also the roughly similar sequencing effort applied to all samples, the cow samples were the most diverse on average (nonpareil diversity = 20.62; nonpareil diversity is in log scale), followed by pigs, then humans (19.4 and 17.6, respectively; Table S2). Nonetheless, the sequencing coverage of these samples was within a suitable range for whole community comparison to identify differentially abundant individual genes/functions or genomes/MAGs (20). NMDS analysis based on MASH distances (8) showed that samples clustered by host type and time, and that the D7 time points did not return to the natural community composition present in the lake water at D0 (Figure S2). Furthermore, ANOSIM analysis of the MASH distances showed that the samples were significantly different (P=0.001) by host type and sampling day (R=0.54 and 0.44, respectively) and ADONIS analysis predicted that these two variables explained (R^2^) 44.0% and 41.3% of the variation in the MASH distances, respectively (P=0.001)

### *Functional annotation for MAGs identified as potential biomarkers and differentially abundant (DA) functions between host fecal metagenomes:*

The 14 MAGs identified as potential markers based on their host sensitivity, abundance, and decay kinetics in the mesocosms were functionally annotated and summarized into KEGG modules using MicrobeAnnotator (Ruiz-P et al. *in review*). There was an average of 2,175 genes in each MAG, of which roughly 48% could be annotated to the KEGG ontology database (2019-04-09 release; (11)). Most of the KEGG modules identified in the MAGs were related to carbon and amino acid catabolism and the reductive pentose phosphate pathway for carbon fixation (Figure S6) suggesting the potential for primary production in addition to fermenting carbon sources derived from the host’s diet. Common fermentation-related genes such as fumarate reductase and succinate dehydrogenase as well as modules for methanogenesis from methanol and methylamine were present, but there was no clear clustering of the MAGs by host type when examining the KEGG modules overall and no modules were clearly unique to a single host type (Figure S6). Thus, DESeq2 differential abundance analysis (13) was used at the gene level to identify specific functions that are DA in the host fecal metagenomic assemblies, which allowed for more comprehensive and sensitive functional comparisons of the host fecal communities (i.e. not restrict the analysis to only the functions that were binned into a MAG). Furthermore, the number of host fecal metagenomic reads that mapped to the collection of high quality fecal MAGs was greater than 2X different between samples (data not shown), which indicated a high likelihood of false-positive results at the MAG level (i.e., finding DA functions by chance due to differences in coverage; (20)). The gene level analysis circumvents this limitation.

Predicted ORFs from the assembled contigs of the host fecal metagenomes were annotated against the KEGG database similarly to the MAGs. There was an average of 107,588 ORFs per assembly, of which ~30% could be annotated with functions other than hypothetical or unknown. The metagenome short reads were mapped against the predicted ORFs to determine their abundances in each metagenome. Of the 2,080 total KEGG functions identified, 177 were significantly DA with P_adj_ < 0.05 and log_2_ fold change (L2FC) > 3 using pairwise comparisons between human, cow, and pig fecal samples. These 177 functions were manually grouped into 39 broader functional categories (Supplementary Data S2) for visualization (Figure S13). The cow fecal assemblies were more abundant in genes related to biofilm formation, starch and sucrose metabolism, and transport of maltose, urea, and putrescine. The pig fecal assemblies were more abundant in genes for amino acid (particularly lysine) degradation, ribosomal proteins and a ribonucleoside-diphosphate reductase (*nrdB*). Human fecal assemblies were enriched in ribose transport, biotin metabolism, and quorum sensing genes. The cow and pig assemblies were more abundant in biosynthesis pathways for amino acids and secondary metabolites, metabolism of cofactors and vitamins, and particularly eight genes for a type IV secretion system (T4SS) related to conjugation (*trbBCDEFGIL*), which were absent in all human fecal assemblies, except for the *trbL* gene, which was only identified in hum3 at low relative abundance (94 reads matching; Supplementary Data S2). Notably, the cow and pig assemblies were also enriched in methanogenesis genes associated with the CO_2_ pathway (21) (*fmdE*; formylmethanofuran dehydrogenase subunit E) and the acetate pathway (*acsD*; acetyl-CoA decarbonylase/synthase complex subunit delta) that were absent in the human assemblies. Human and cow assemblies had more genes related to zinc transport, lipid metabolism, and the T6SS secreted protein *vgrG* compared to pigs, whereas human and pig assemblies were more enriched for nitrogen, ascorbate and aldarate metabolism compared to cows (Figure S13). Despite being from a strictly anaerobic environment, the cow and human gut samples were more abundant for catalase, which may indicate that these guts are more prone to aerobiosis e.g., from rapid biomass growth or infected epithelial tissues in the GI tract (22) compared to the pig gut.

Of the 177 significantly DA KEGG genes identified by DESeq2, 137 were also present in at least one of the putative biomarker MAGs described above. An average of 27 ±9, 23 ±18, and 10 ±9 DA genes were found in the human, cow, and pig putative biomarker MAGs, respectively (Supplementary Data S4). However only seven of these were found in all three human MAGs and none were found in all five cow or six pig MAGs indicating that the host-specific functions are not shared consistently among the corresponding, host-specific MAGs. Alternatively, incomplete MAGs could account for these findings, i.e., these MAGs ranged between 70-95% in their completeness estimate (Supplementary Data S1). Furthermore, two of the pig MAGs (pig7_006_Acetobacteroides_C and pig8_10_Acetobacteroides_C) had only two DA KEGG genes, one of which was a glycine dehydrogenase subunit I gene (present in 4/6 MAGs). The 7 DA KEGGs that were found in all 3 human MAGs included a lactaldehyde reductase, hydroxylamine reductase, transaldolase, and several genes related to vitamin B12 synthesis. The most common KEGG gene present in the cow MAGs was pullulananse, which was present in 4 of 5 cow MAGs (Supplementary Data S4). Together, these results reinforce that most of the putative host-specific MAGs are robust targets because they contain genes that were also DA at the assembly level.

### *Decay of Lake Lanier (LL) MAGs:*

### The mesocosm metagenomic shorts reads were searched against a collection of 477 Lake Lanier (LL) MAGs (i.e., the native microbes present in the source lake water representing six years of surface water samples from Lake Lanier; (23)) to assess MAG abundance dynamics over the incubation time. Of the 477 LL MAGs, 59 (average 22 MAGs per mesocosm) and 139 were detected in the host fecal (Figure S9) and negative control (Figure S11) mesocosm samples, respectively. The different host fecal inocula (and their associated communities) did not have a consistent effect on the abundance of the native LL MAGs over the time-series (Figure S9). Overall, the LL MAGs showed much lower abundance than the host fecal MAGs in the mesocosms (<0.7%) and varied, more or less randomly, in their relative abundances with time, consistent with the assumption that they represent autochthonous taxa present in the source water used in the incubations.

### *Decay kinetics of human mtDNA qPCR assay and reference genome:*

### Overall, the two *Bacteroides* markers (RumBac and HF183 in H3) had more similar decay characteristics compared to the HUMmt marker in the dialysis bag mesocosms, and the 2-log reduction time (t_99_) was slightly longer for HUMmt (10.6 d) compared to both HF183 (6.8 d) and RumBac (7.5 d) in H3 (Table S5). These results indicated that the mtDNA is able to persist longer in the extraenteric environment which may be due to the added layer of protection offered by the mitochondrial membrane in addition to the cytoplasmic membrane of eukaryotic cells.

Since it was not possible to estimate the human mtGenome copy number per cell, the HUMmt qPCR assay estimated counts per mL were compared to the relative abundance of a reference human mtGenome (Table 1) expressed as sequencing depth per metagenome size (in Gbp) and a weak correlation (R^2^=0.29) was observed (Figure S17). Although these results are clearly based on a limited sample size, the human-specific assays were consistent with known limitations for interpreting qPCR results, namely, host-sensitivity issues for HF183 and lack of absolute abundance estimates for HUMmt.

### *Comparisons of putative MAG biomarkers against traditional FIB and MST assays:*

We compared traditional qPCR-based abundances of common MST markers to metagenome-based methods. We estimated total cell densities from GenBac16S qPCR counts to estimate absolute abundances (i.e., cells/mL) of the corresponding MST reference genome in the metagenomes. The ruminant-specific *Bacteroidetes* 16S assay, RumBac, consistently over-estimated the abundance compared to the metagenome-based methods. This could be due, at least partly, to the small amplicon size (118 bp), which is typical for most qPCR assays and could be detectable even in highly degraded DNA from dead cells (24). In contrast to RumBac results, the human-specific HF183 assay was poorly correlated and consistently under-estimated the abundance of *Bacteroides* compared to metagenome-based methods. This was most obvious in the H1 mesocosms where qPCR-based estimates reported < 7 *Bacteroide*s cells/mL whereas metagenome-based estimates showed 0.5-3x10^6^ *Bacteroides* cells/mL in D0, D1 and D4 samples. Our further investigation revealed several mismatches between the forward primer and the assembled contigs that carried the target gene of the HF183 assay (Figure S15). Neither contigs were binned into fecal MAGs, which is not surprising because 16S-carrying contigs are often problematic for the abundance-based binning methods used here. These findings most likely accounted for the low HF183 qPCR-based *Bacteroides* abundance estimates compared to the metagenome-based estimates (i.e., less or no exponential doubling because PCR amplification is only happening at the reverse primer). It is unlikely that this discrepancy is the result of PCR amplification inhibition because DNA samples were diluted in water before running qPCR. Furthermore, negative control samples also indicated no added 16S counts to the cell density estimates. The amplification efficiency for the HF183 and GenBac16S assays were relatively low (87.9% and 81.2%, respectively; Table S4). However, in the case of HF183 in H1, qPCR estimates were 5 to 6 orders of magnitude less than metagenomic estimates, thus this difference is too large to come from an 88% amplification efficiency alone and, instead, is likely the result of primer mismatches mentioned in the main text or a combination of this and other explanations. The latter conclusion is further supported by the fact that HF183 estimates in H3 were more similar to metagenomic estimates (Figure 3A) and thus, a systematic under-estimation of the qPCR reaction is unlikely to explain the discrepancy observed. A lower % efficiency is expected for the GenBac16S assay, which uses degenerate primers and thus is less accurate at providing exact estimates for 16S copy numbers. Nevertheless, microbial load biases are a significant issue for evaluating compositional changes in metagenomic datasets (25, 26) and using the GenBac16S assay allows to control and correct for relative changes in total cell density (Figure 1D). Furthermore, the tendency for this assay to under-estimate cell density is presumably systematic across all samples and thus, it is still useful for converting metagenomic-based relative abundance to absolute abundance estimates. Indeed, using the GenBac16S assay represents one of the best-known methods available for improving the quantitative capabilities of metagenomic data (27, 28). Although the absolute abundance estimates provided here may not be highly accurate, they do provide preliminary evidence that quantitative metagenomic approaches based on GenBac16S and similar assays can produce results that are similar to qPCR methods (especially in the case of RumBac) and thus, expand the toolbox of microbial source tracking. However, these results should be confirmed with further testing against samples from other habitats and conditions.

Targeting a single marker (whether it be a whole genome or qPCR assay) for MST can still be inadequate for water quality monitoring because of the high inter-person variability observed in the human gut. For example, neither the HF183 assay nor the *B. dorei* reference genome were detected in any of the H2 mesocosms (or the hum2 feces), thus these water samples, although highly polluted with human feces in our experimental set up, would have been considered safe for public health if using only this single *Bacteroides* marker. The importance of inter-person variability/specificity was also evident when looking at the human fecal MAG abundances in the mesocosms (Figure S7A). The most abundant MAGs in each human mesocosm corresponded to the human fecal sample that was used as an inoculum for that mesocosm (e.g., hum1 MAGs were the most abundant in H1 mesocosms). Typically, the most abundant MAG in each mesocosm was ~10X more abundant than MAGs from the other two human fecal samples on average. Although both metagenomics and qPCR have unique benefits and limitations, the results presented here suggest that using only a single marker to assess human fecal pollution was inadequate for either method. This result is not surprising considering previous studies of human gut communities that have revealed extensive diversity among individuals (29, 30) and intra-person temporal variability within the human gut microbiome, which suggests that no core taxa exist that are abundant in each human microbiome (31). Although the relative abundance and taxa can vary over time, significant functional redundancy has been observed previously (32, 33) as well as in our human MAGs (Figure S13). Therefore, if the goal is to use a single or a few genes as biomarkers then functional genes as opposed to the 16S rRNA gene or individual taxa (e.g., MAGs) may be a more robust strategy due to the high prevalence of some gene functions among individuals of the same host type, presumably driven by the host-specific gut physiology. Metagenomics, as shown in this study, can be useful in identifying these genes to be used as novel targets for more robust qPCR assays.

### *Bottle effect in mesocosms on D7:*

There was evidence of a bottle effect in the dialysis bag mesocosms that became apparent at the D7 time point. Nonpareil (9) results showed an increase in overall community coverage (and thus a decrease in community diversity) that reached ~76% on D7 for all samples including the negative controls over time in the dialysis bags (Table S1). Furthermore, the average genome size as determined by MicrobeCensus (10) also showed an increase with time (Figure S1). The high coverage in the D7 mesocosm samples indicated that it was possible to recover some MAGs from these samples (as opposed to the earlier time points, which were characterized by too high diversity based on Nonpareil to expect good assemblies or MAGs). Contig binning from the D7 mesocosm metagenomes resulted in 39 high quality MAGs that dereplicated into 17 genomospecies at 95% ANI. These MAGs appeared to be highly different from those assembled from the host fecal communities based on pairwise AAI comparison (Figure S12) that showed none of the D7 MAGs formed a genomospecies (ANI >95%) with any of the host fecal or Lake Lanier (LL) MAGs. Taxonomic classification of these MAGs using MiGA and the TypeMAT/NCBI database showed that the closest relative to three of these MAGs (AAI ~47%) was *Methylobacterium platani* and the closest relative to four other MAGs (AAI ~66%) was *Cellvibrio japonicus*; two additional MAGs were classified in the family *Cytophagaceae* (Supplementary Data S1). Therefore, these D7 MAGs were, in general, more related to each other than the host MAGs. Furthermore, none of the D7 MAGs were classified as class *Bacteroidia* or *Clostridia*; the most common taxa among the fecal MAGs. When all the MAGs were clustered by phenotype using Traitar v1.0.4 (18), the D7 MAGs formed a distinct cluster from the rest of the host fecal MAGs and all D7 MAGs were classified as aerobes with oxidase and catalase (Figure S5). When looking at the abundance of the D7 MAGs over time in the mesocosm metagenomes, they were not detectable in any of the D0 or D1 time points and became detectable in only a few metagenomes by D4 (Figure S10). However, on D7, they increased to 30-50% of the total community in some cases, and accordingly the majority of the metagenome short reads from D7 samples also mapped to these MAGs (Figure S3). The LL MAGs in the negative control dialysis bags tended to decay over time and were mostly undetectable by D7 when the D7 MAGs started to increase (Figure S11). These results suggest that the bottle effect in the D7 samples was consistent across all mesocosms and was not very substantial before D4.

The results of the D7 MAGs were further confirmed when comparing the relative abundances of the genes recovered in the D7 assemblies against the genes of the host fecal assemblies using DESeq2. Out of 2,906 total KEGG functions detected, 582 were significantly DA with P_adj_ < 0.05 and L2FC > 6 that were manually grouped into broader functional categories as described in (Supplementary Data S3). There were many more significantly DA genes when comparing the D7 samples to all of the animal host fecal metagenomes, and the differences were much greater and more distinct than in the animal host only comparisons described above. The D7 samples were particularly enriched for genes related to aerobic processes (Figure S14 and Supplementary Data S3) such as cytochromes and sulfide oxidation genes, consistent with results from the Traitar analysis (Figure S5). Furthermore, the D7 samples were enriched for genes related to photosynthesis and porphyrin and chlorophyll metabolism and biofilm formation. In contrast, the host fecal metagenomes were more enriched in genes for anaerobic processes like methanogenesis, acetylaldehyde/alcohol dehydrogenase, sulfur and fumarate reductases, nitrogenase (NifH), glycerol dehydrogenase, butyrate kinase, and phosphate butyryltransferase as well as ABC transporters for sugars and amino acid and cobalamin biosynthesis genes. These results suggested that the populations arising by D7 are likely “weed” species from the rare biosphere of the lake water used in the mesocosms that were able to form biofilms on the material of the dialysis bags and perform aerobic metabolism.

Furthermore, the bottle effect was not apparent until D7; that is, after the point when most of the fecal organisms apparently had died off around D4. Therefore, this bottle effect is not expected to have a major impact on patterns reported here (e.g., host MAG dynamics) or on our conclusions because the latter were primarily based on MAG abundance dynamics during the first four days. This is consistent with other similar studies such as Ahmed et al. 2018, in which sewage OTUs were not detected after four days, and the study found that the mesocosms did not return to the initial community composition even after 50 days (34). Consistently, we did not see a return to starting community within the duration of our experiments (7 days). Mattioli et al. 2018 also reported bottle effects from dialysis bag mesocosms (e.g., lower nutrient concentrations and higher chlorophyll a) that did not arise until after five days post-perturbation (35). These results confirm that delayed bag effects are a common limitation of the dialysis bag mesocosm method and that experiments carried out for longer than 5 days should anticipate further testing to distinguish effects of the experimental design and genuine microbiome changes. Further, it would be important to test these findings with field samples from recent pollution events to further corroborate the abovementioned conclusions [the possibility that our mesocosms have gone anaerobic, and the likely effects of this on MAG abundance dynamics, is provided below].

*Oxygen limitation in the dialysis bag mesocosms:*

It is also possible that the mesocosms became oxygen-limiting, especially during the first 4 days of incubation. Theoretical models for dissolved O_2_ and biological oxygen demand (BOD) over time in the mesocosms (assuming BOD for feces is ~36 mg O_2_/mL and dissolved O_2_ at D0 is 9 mg/mL) show that dissolved O_2_ in the aquarium tanks is rapidly consumed by D1 and doesn’t start to increase again until around D5 (Figure S18). This model coincides well with the inferred strict anaerobic physiology (Figure S5) of the MAGs that peaked in abundance over time in the mesocosm metagenomes. On D4, most of the fecal MAGs have died off (i.e. have zero TAD80), nor have the aerobic D7 MAGs have started to grow substantially (Figures S7, S3, and S10). Then at D5 when O2 levels start to increase, the D7 MAGs are able to aerobically metabolize the remaining BOD in the mesocosms and result in the high abundance of these MAGs seen on D7 (Figure S10). Thus, this scenario reflects a potential outcome for surface waters receiving runoff or sewage rich in BOD; hence, our mesocosm approach was realistic from this perspective as well.

**Table S1:** Total community covered by our sequencing effort as determined by Nonpareil v3.0. Averages from three biological replicates and standard deviation from the average are shown.

| Day | Cow | Pig | Human | Neg Control |
| --- | --- | --- | --- | --- |
| **0** | 26.9 ± 2.4 | 49.5 ± 13.0 | 79.6 ± 3.4 | 46.4 |
| **1** | 27.4 ± 6.8 | 55.0 ± 3.0 | 80.9 ± 1.9 | 53.6 |
| **4** | 54.9 ± 4.0 | 59.2 ± 2.3 | 74.2 ± 10.9 | 70.8 |
| **7** | 79.6 ± 5.5 | 72.5 ± 9.5 | 77.7 ± 2.4 | 76.0 |

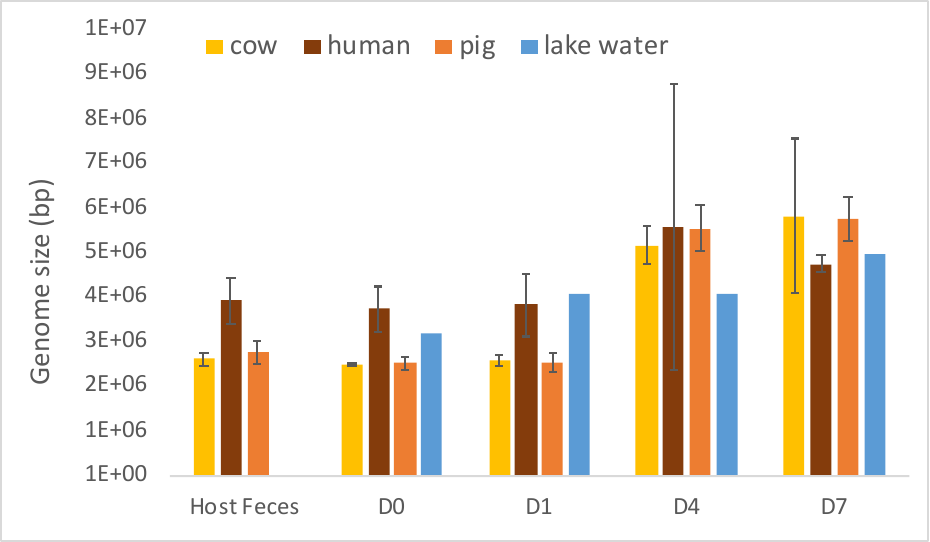

**Figure S1: Average genome size in the host feces, mesocosm and lake water negative control metagenomes as determined by MicrobeCensus.** Error bars are standard deviations from the average based on three replicates except for negative controls which did not have replicates.

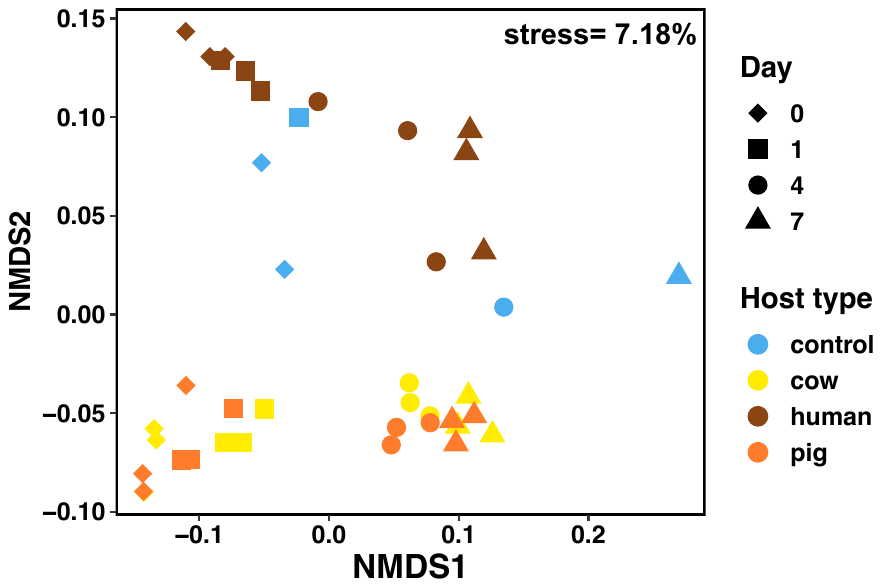

**Figure S2: Similarity among the sequenced communities during the mesocosm incubations.** Graph represents the non-metric multidimensional scaling (NMDS; 4 dimensions) of the whole-community MASH distances (i.e., overall kmer similarity of microbial communities). Each point represents a metagenome sample and samples from the same host type (or negative control) are denoted by the same color. Samples that are more similar are grouped closer together.

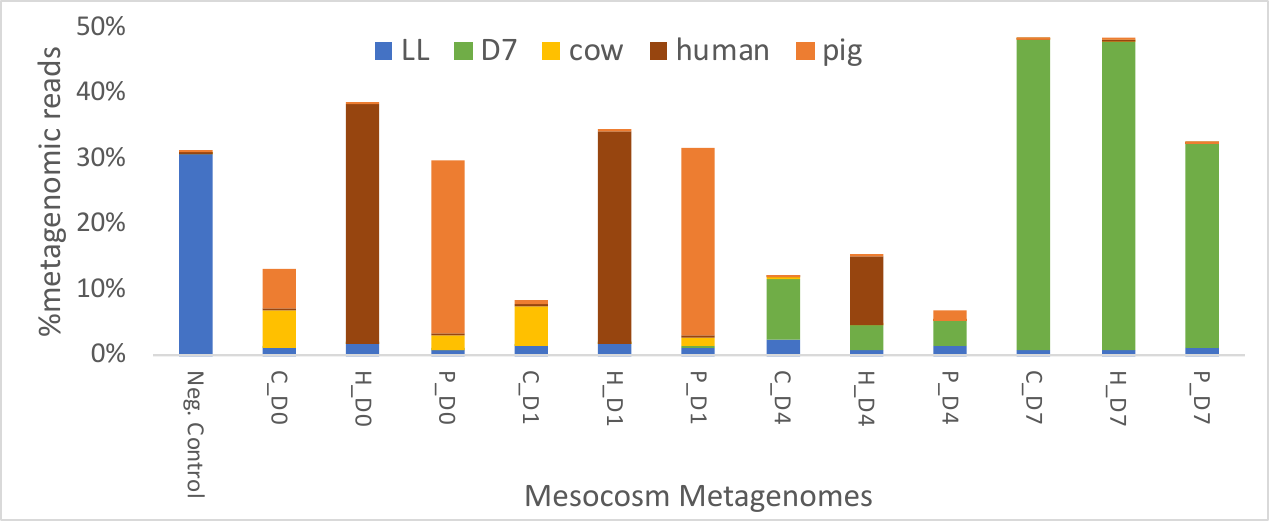

**Figure S3: Fraction of the mesocosm microbial communities represented by the LL, D7, and fecal MAGs**. The graph shows the average number of mesocosm metagenome short reads that matched to any of the 477 Lake Lanier (LL), 17 D7, 17 cow, 13 human, and 49 pig fecal MAGs that were used in this study. Each column represents the mesocosm metagenomes by host type through time (e.g., C_D0 is the average of all 3 cow biological replicate mesocosm metagenomes at D0). Read counts were normalized by the total number of reads in each mesocosm metagenome (expressed as % of total number of short reads).

**Table S2: Metagenome sample, trimming and Nonpareil coverage and diversity information for dialysis bag mesocosm and host fecal metagenomes sequenced in this study.** Results are reported for reads after quality trimming and removal of host DNA with bmtagger (for fecal samples only).

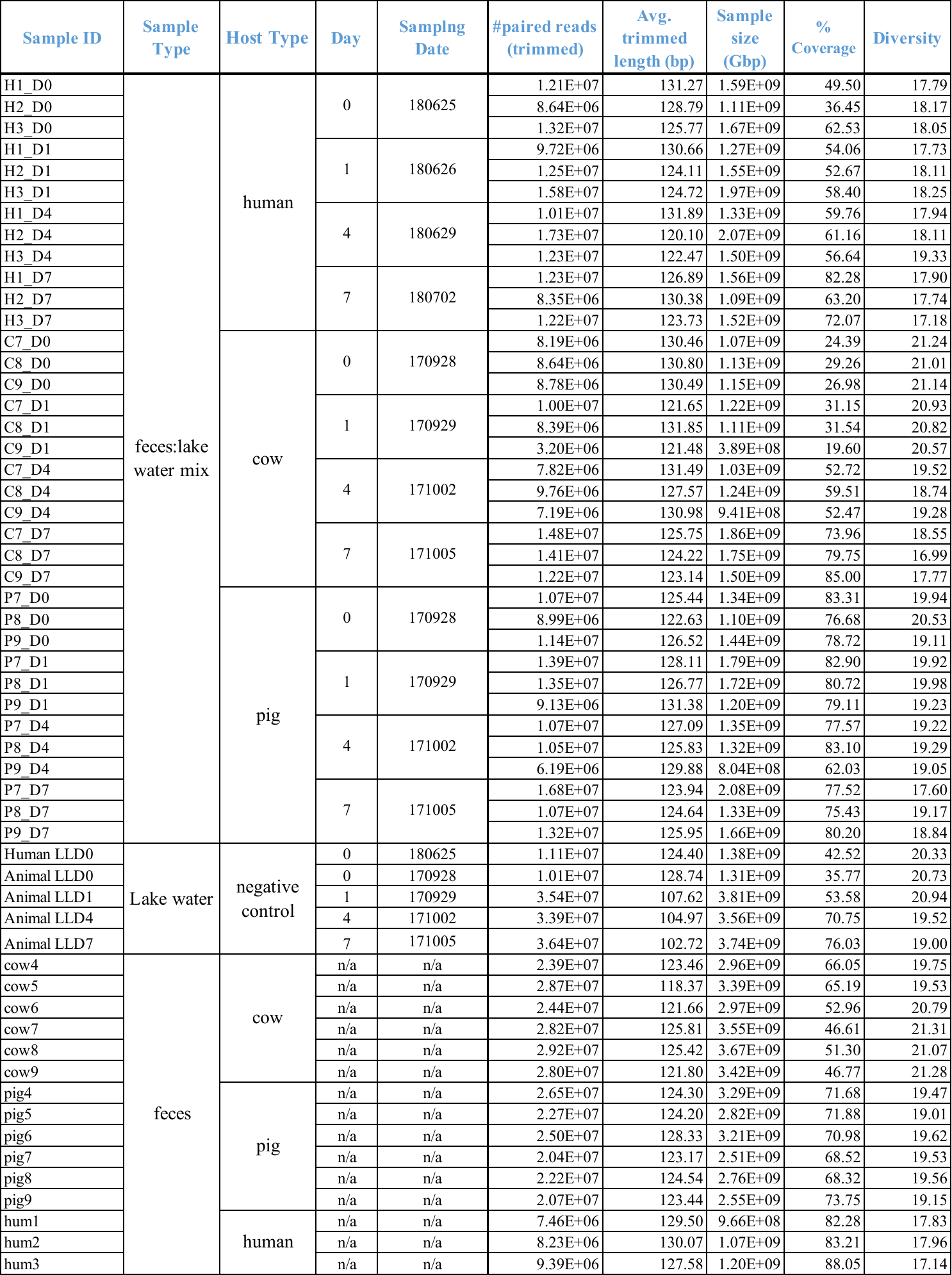

**Table S3: Assembly information for host fecal and D7 metagenomes.**

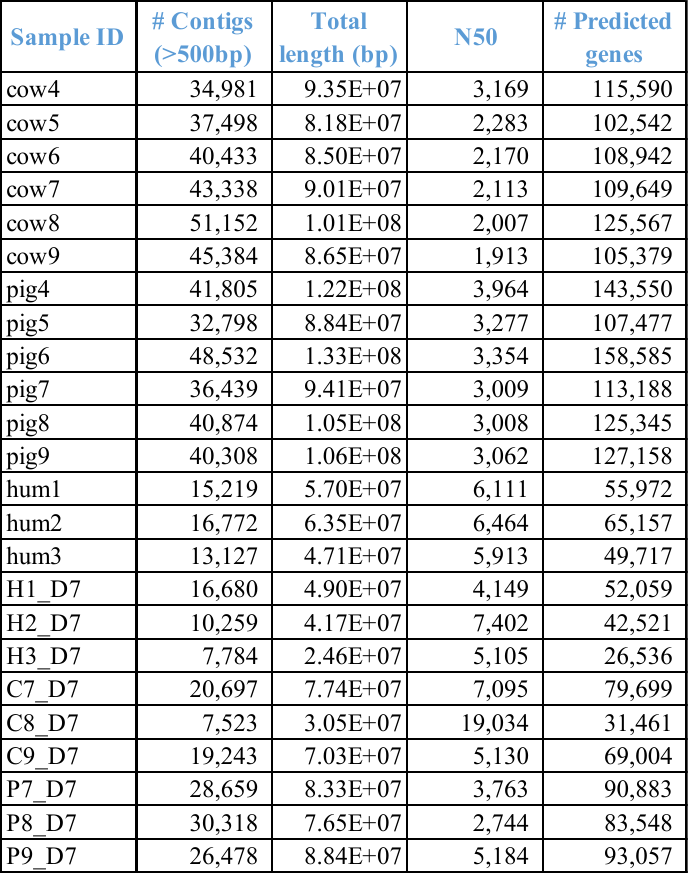

**Table S4: qPCR assay reaction details and performance.**

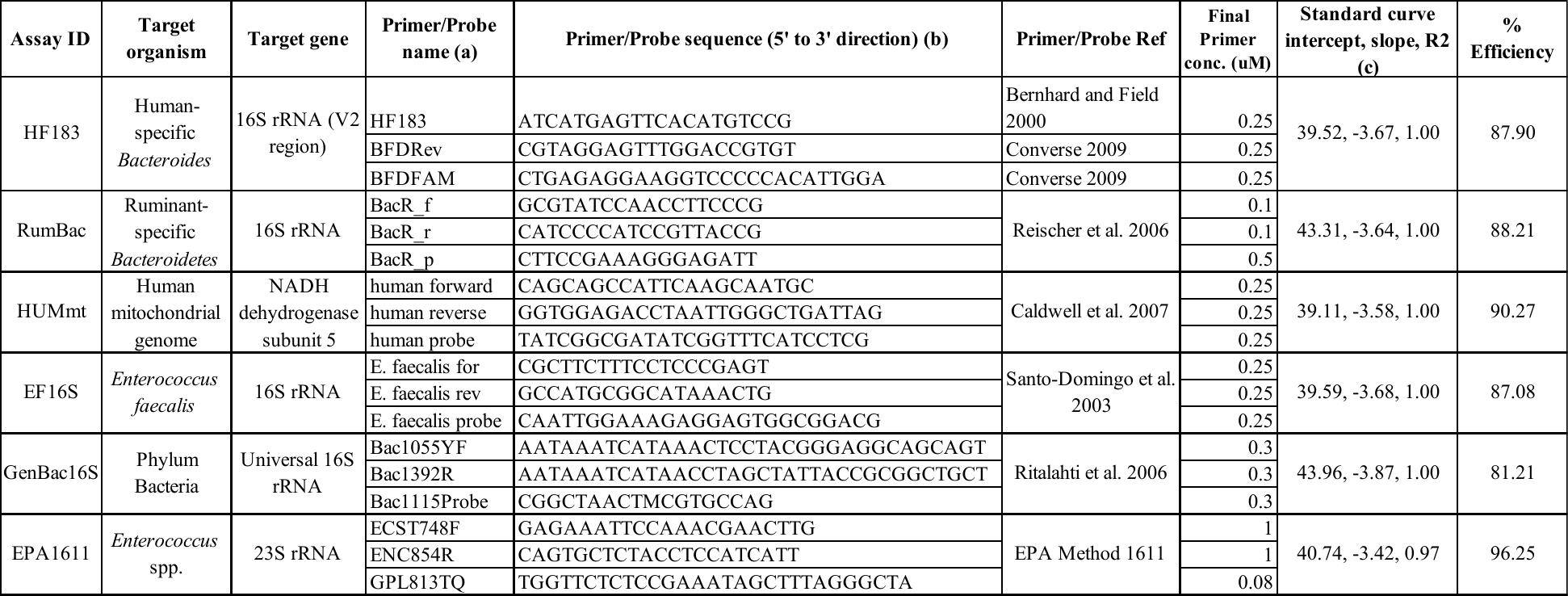

**(a)** Primers and probes are listed in the following order: forward, reverse, hydrolysis probe.

**(b)** All of the hydrolysis probes were labeled at the 5' end with the reporter dye FAM (6-carboxyfluorescein) and at the 3' end with a non-fluorescent Iowa black quencher (BHQ) with a minor groove binding moiety. Except the EPA1611 assay which used a TAMRA quencher dye as described in the EPA Method 1611.

**(c)** Reported as the average for all plates ran EXCEPT for EPA1611, in which results for the composite curve used to calculate the average number of target sequences in calibrator cells are reported.

**Table S5: First order decay characteristics for MAGs and MST markers in the dialysis bag mesocosm metagenomes**. Only biomarkers detected in at least 3 timepoints were used. MAGs identified as putative biomarkers are in bold. C0 is the absolute concentration at 0 in cells/mL for all unless noted otherwise in parentheses in the second column.

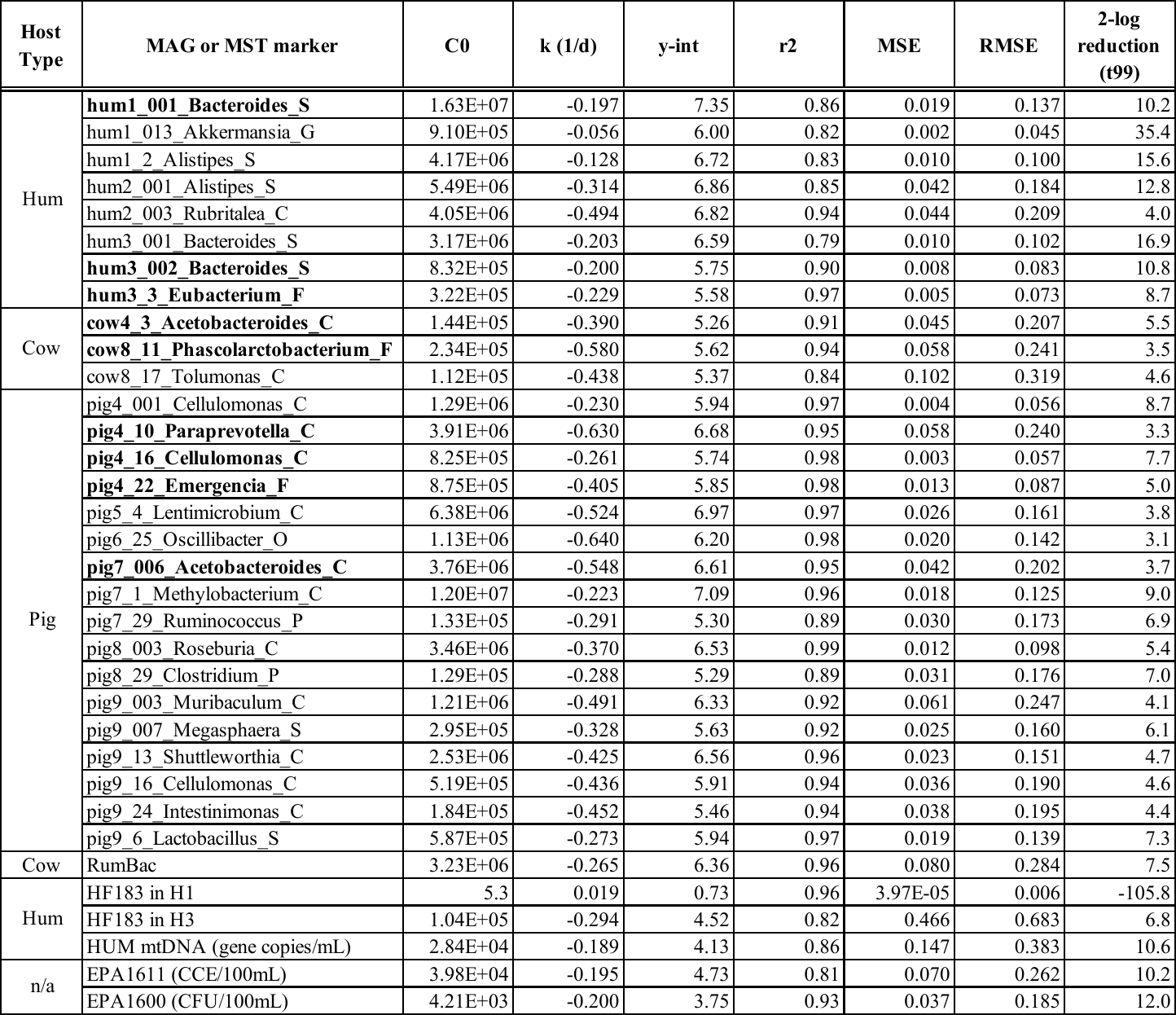

**Table S6: Detection of MST qPCR markers in feces (inocula material) or un-inoculated lake water (negative control).** Values are reported as average number of copies per mg or mL for fecal and lake water samples, respectively. ND = Not Detected, which indicates that sample did not return a Ct value for all biological replicates. DNQ = Detectable but not Quantifiable, which indicates that sample returned a Ct value that was higher than the lowest concentration in the standard curve.

| **Marker** | **Pig feces** | **Cow feces** | **Human feces** | **Lake water** |
| --- | --- | --- | --- | --- |
| RumBac | ND | 4.1x10^6^ ±6.1x10^5^ | ND | ND |
| HF183 | ND | ND | 8.9x10^4^ ±1740 | DNQ |
| HUMmt | ND | ND | 1645 ±316 | DNQ |
| EF16S | 141 ± 51 | 153 ± 59 | ND | ND |

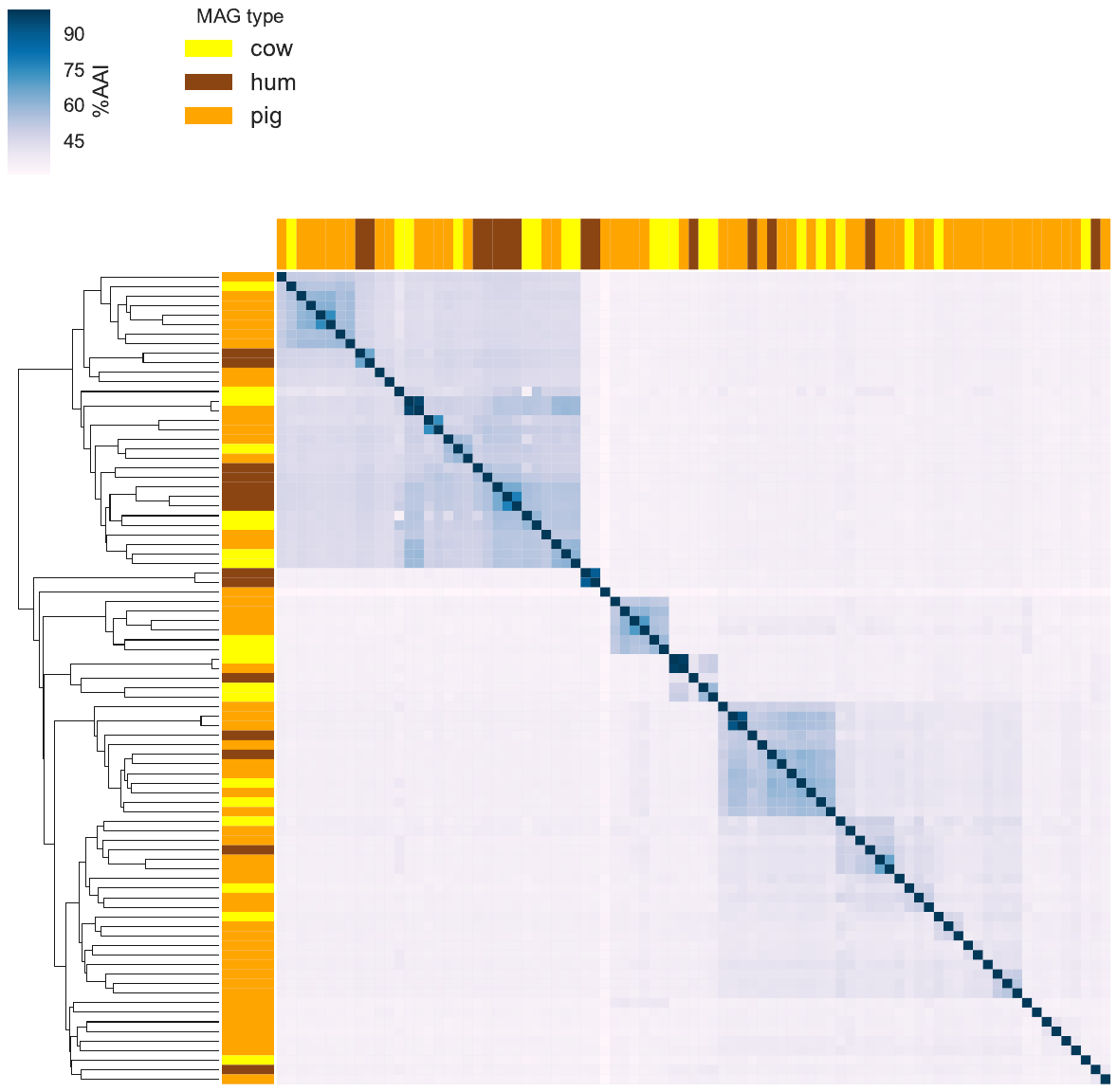

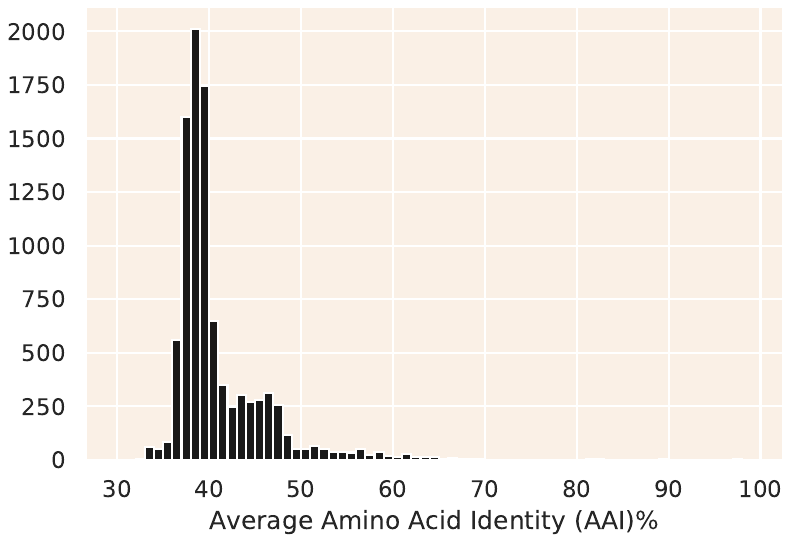

**Figure S4: Genetic relatedness among the fecal MAGs recovered by our study.** Heatmap comparing average amino acid (%AAI) of the MAGs assembled from pig, cow, and human fecal metagenomes (top). Histogram of % AAI values (bottom).

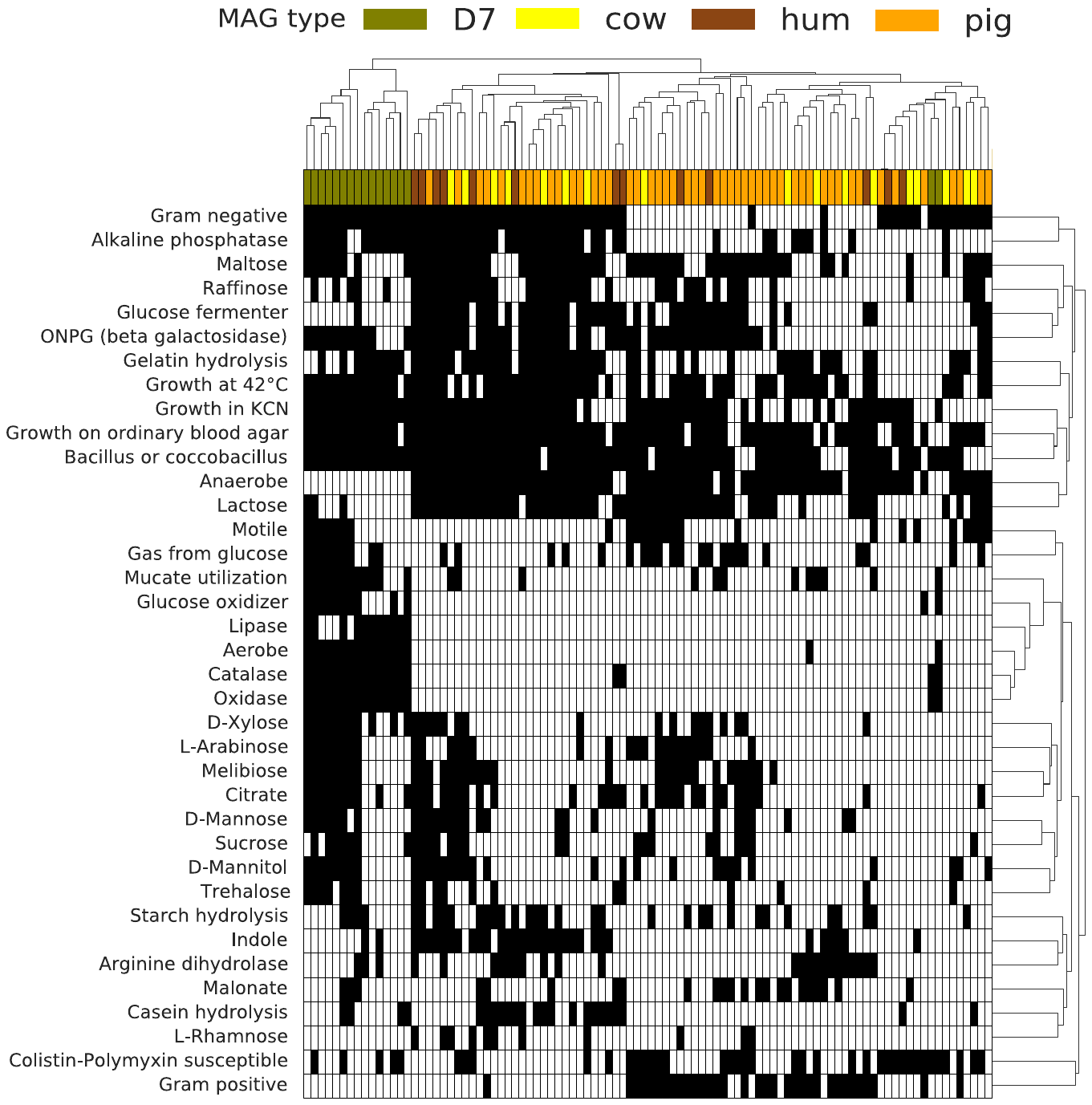

**Figure S5: Heatmap of the presence (black) and absence (white) for different phenotypes of all 96 MAGs as determined by Traitar**.

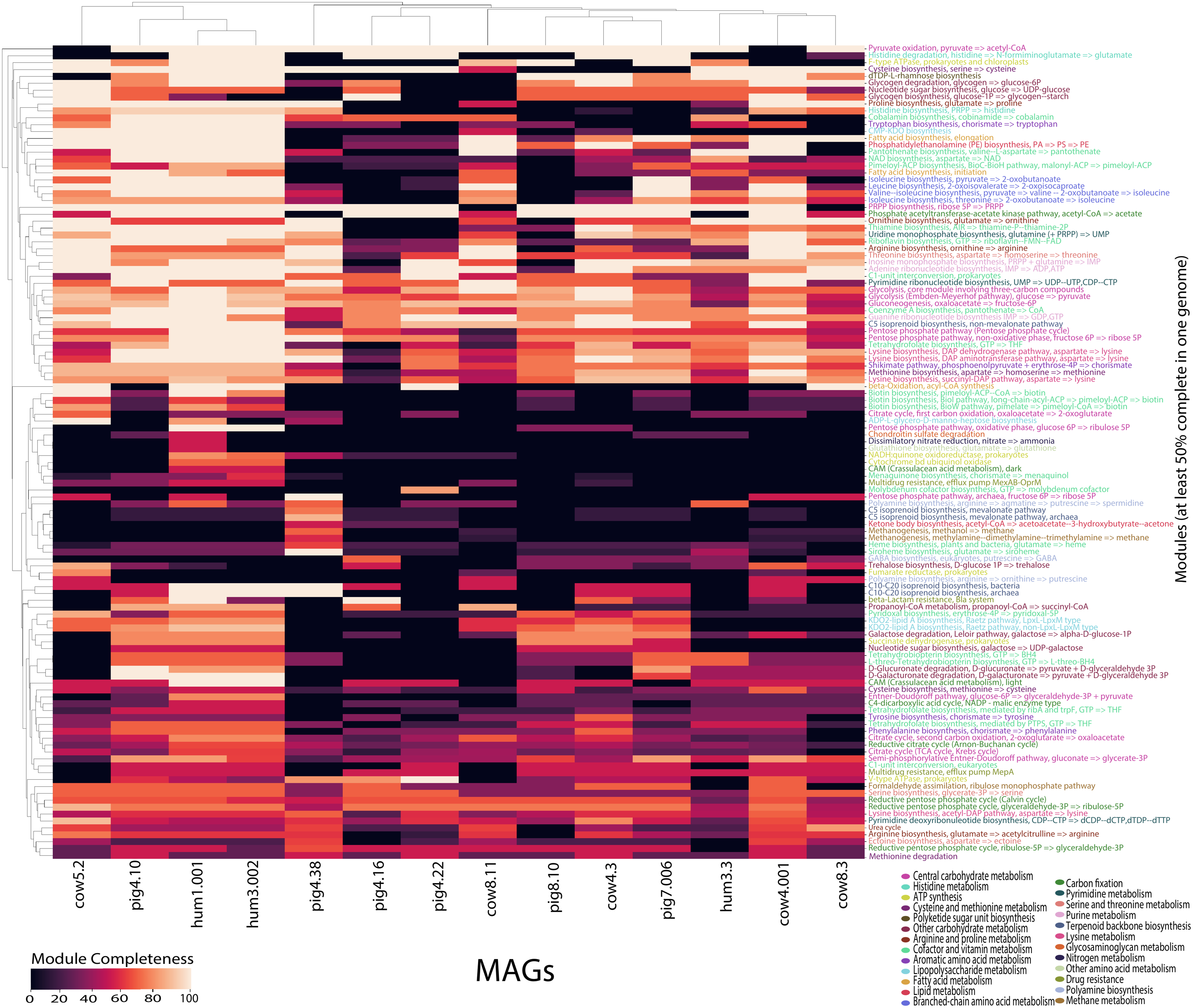

**Figure S6:** **KEGG modules present in host MAGs identified as potential biomarkers**. Only the MAGs identified as potential biomarkers (see Figures S7 and S8) are shown here for visualization purposes.

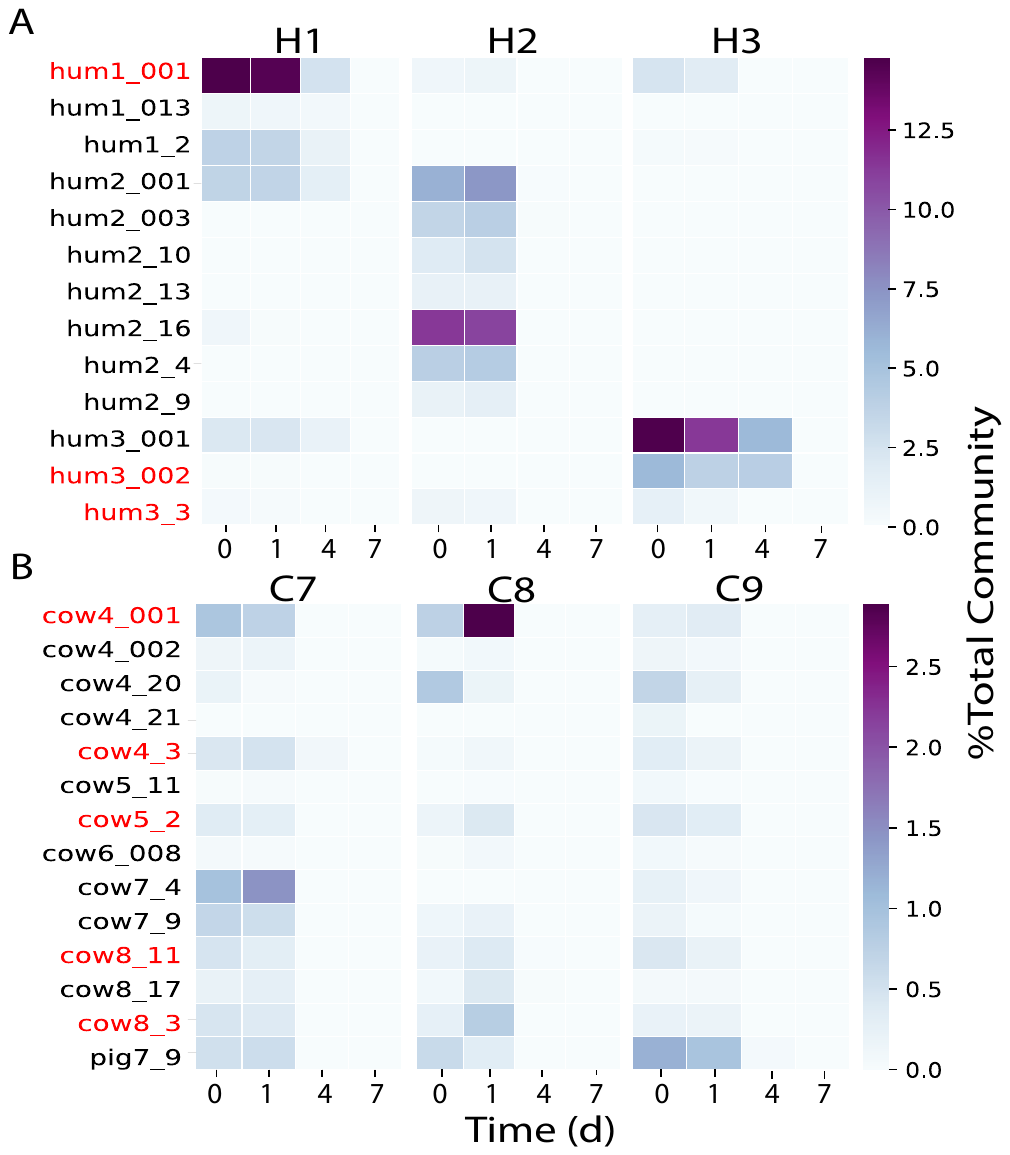

**Figure S7: Decay kinetics of all host fecal MAGs (rows) that could be detected in the (A) human and (B) cow mesocosms (columns).** Note that no non-target host MAG was detected in any of the human mesocosms; the single pig MAG shown was clustered into a single genomospecies at >95% ANI with another cow MAG that is not shown here because the pig MAG was of higher quality and was used in all downstream analyses instead of the MAG obtained from the cow metagenome. H1, H2, H3 and C7, C8, C9 are the different biological replicate mesocosm metagenomes for each host type. Abundances are reported as % of total bacterial community (i.e. TAD80 divided by average genome sequencing depth; details in the main text). The MAGs identified as potential MST biomarkers have red labels. The MiGA TypeMat/NCBI taxonomic identifications appending the MAG names as described in the main text are not included here due to space limitations (see Supplementary Data S1).

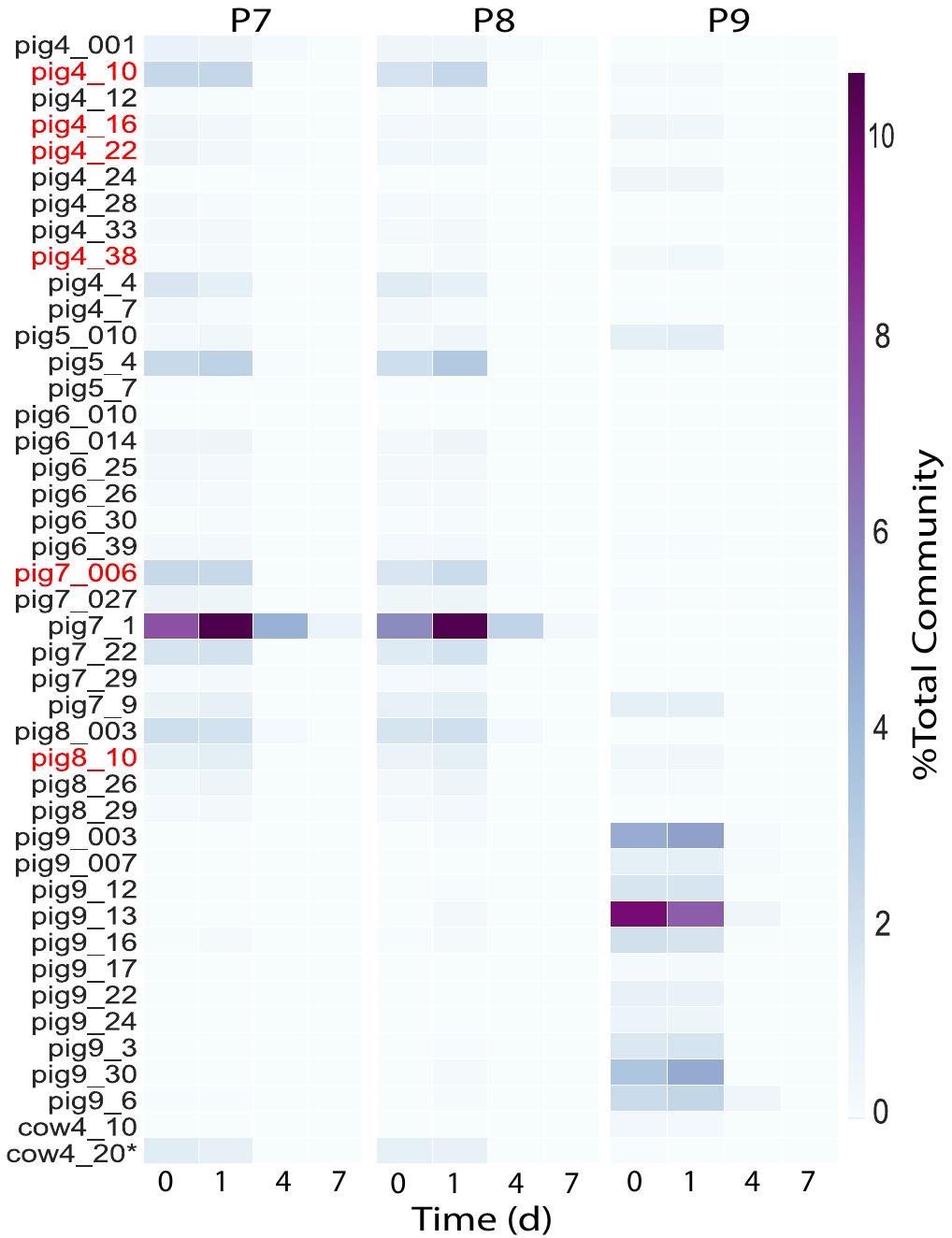

**Figure S8: Decay kinetics of all host fecal MAGs (rows) that could be detected in the pig mesocosms (columns).** The MiGA TypeMat/NCBI taxonomic identifications of the MAG names (Supplementary Data S1) are not included due to space limitations. Cow4_20 clustered into a single genomospecies at >95% ANI with a pig MAG that is not shown here because MAG cow4_20 was of higher quality and was used in all downstream analyses instead of the MAG obtained from the pig metagenome. P7, P8, and P9 are the different biological replicate mesocosm metagenomes. Abundances are reported as % of total bacterial community (i.e. TAD80 divided by average genome sequencing depth; details in the main text). The MAGs identified as potential MST biomarkers have red labels.

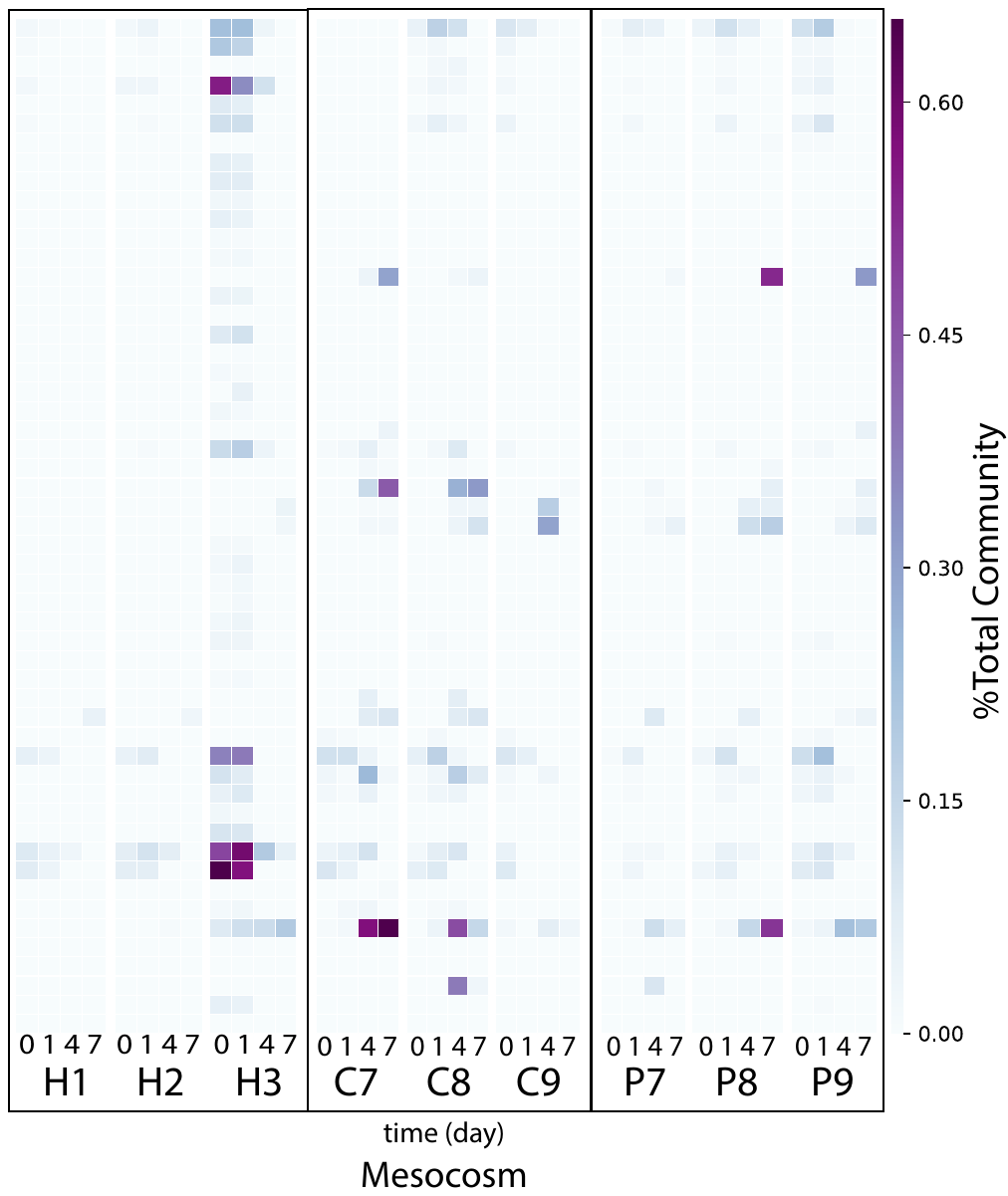

**Figure S9: Abundance kinetics of Lake Lanier (LL) MAGs in the fecal mesocosm samples over time.** The collection of 477 LL reference MAGs from Rodriguez-R et al. 2020 (23) was searched against the time-series mesocosm metagenomes and the relative abundance for 59 MAGs that were detectable are shown as individual rows in the heatmap. Each column is a mesocosm metagenomic dataset and the 0, 1, 4, or 7 refers to the sample time in days while the H, C, or P refers the human, cow or pig biological replicate mesocosm (e.g. the “0” column above H1 is hum1 mesocosm at day 0). Abundance is expressed as % of total bacterial community.

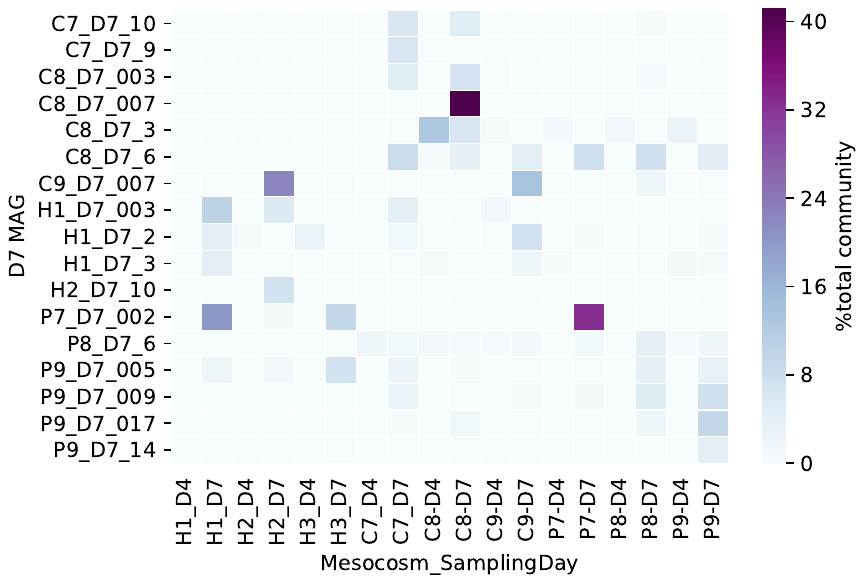

**Figure S10: Abundance kinetics of Day 7 (D7) MAGs in the fecal mesocosm samples over time.** The collection of 17 D7 MAGs assembled from the D7 mesocosm metagenomes was searched against all time-series mesocosm metagenomes and their relative abundances (rows) are shown. Each column is a mesocosm metagenome and only the D4 and D7 samples are shown because no D7 MAGs were detectable in any of the earlier time points. Naming format: number refers to the sample time in days while the H, C, or P refers the human, cow or pig biological replicate mesocosm. Abundance is expressed as % of total bacterial community.

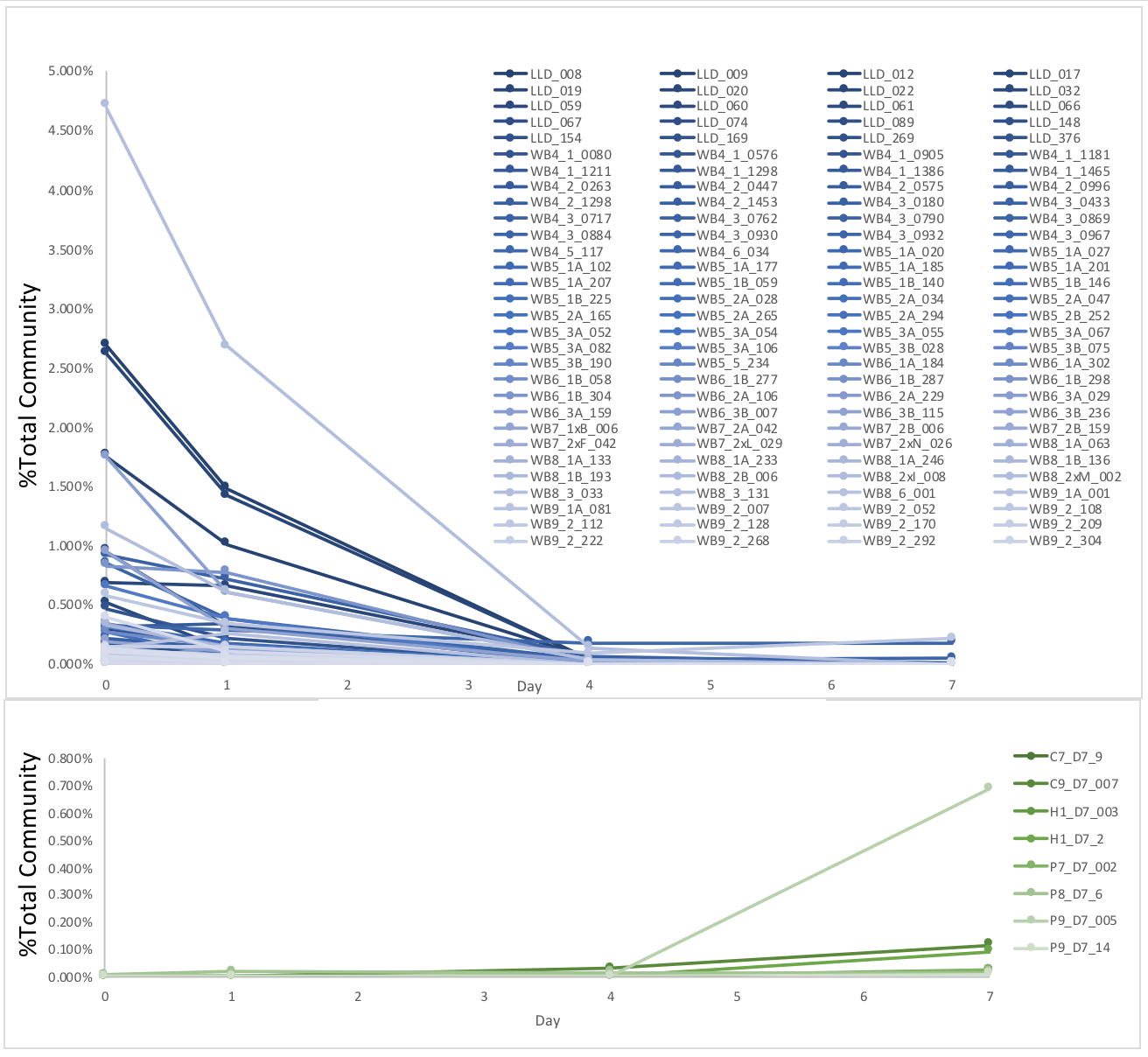

**Figure S11: Decay kinetics of LL and D7 MAGs in the uninoculated lake water negative control dialysis bags.** None of the host fecal MAGs nor the MST reference genomes were detected in any of the negative control metagenomes. Abundance is reported as the percent of the total bacterial community. **(Top)** Decay of the 139 LL MAGs that could be detected in any of the negative control mesocosms from the 477 LL MAG collection (23). (**Bottom**) Only 8 of the 17 D7 MAGs could be detected in the D7 negative control sample. No D7 MAGs were detected in any of the earlier time points.

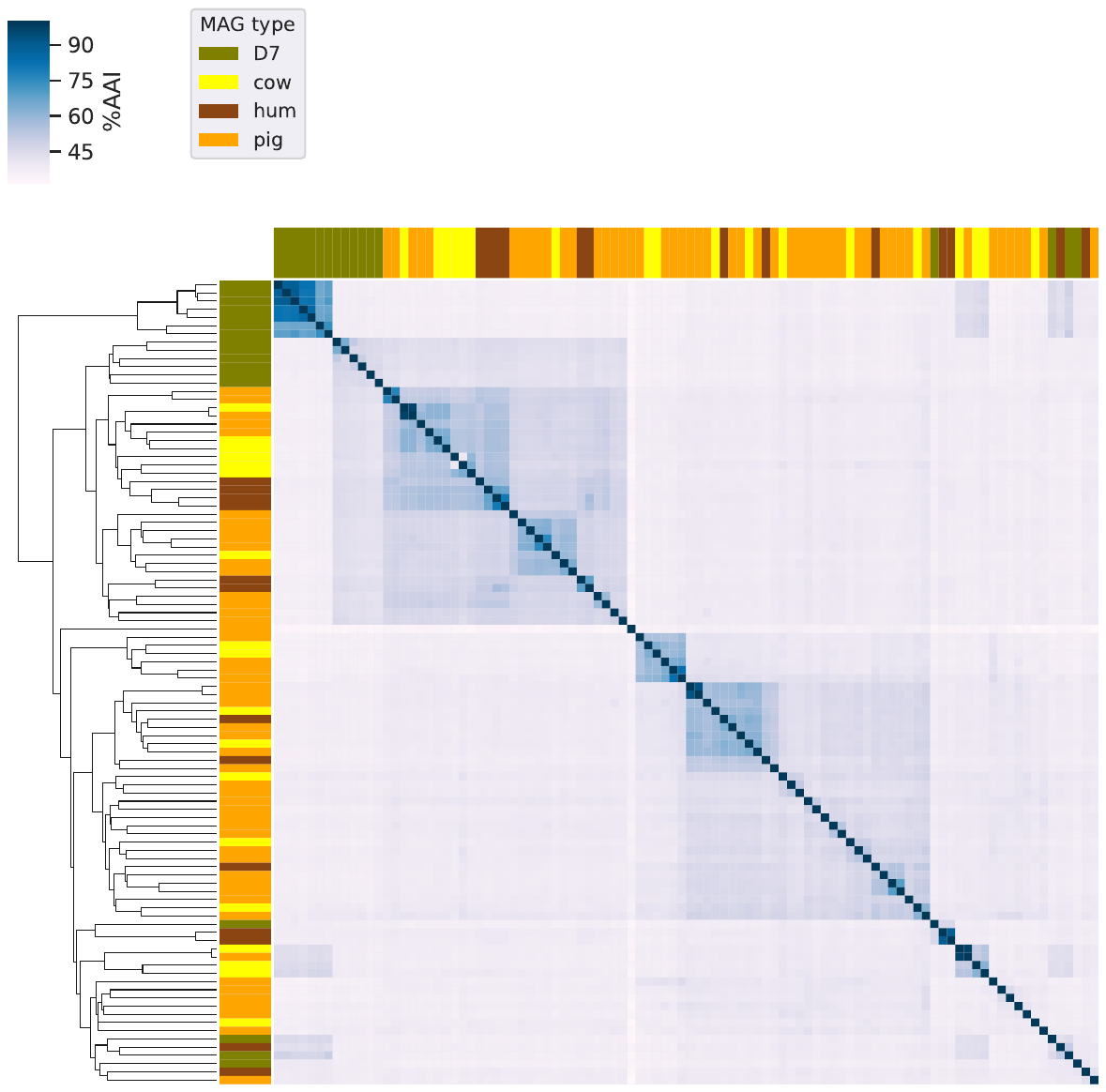

**Figure S12: Genetic relatedness among the host fecal and D7 MAG recovered by our study.** Heatmap comparing average amino acid (%AAI) of the MAGs assembled from pig, cow, and human fecal and D7 mesocosm metagenomes (see figure key on the top).

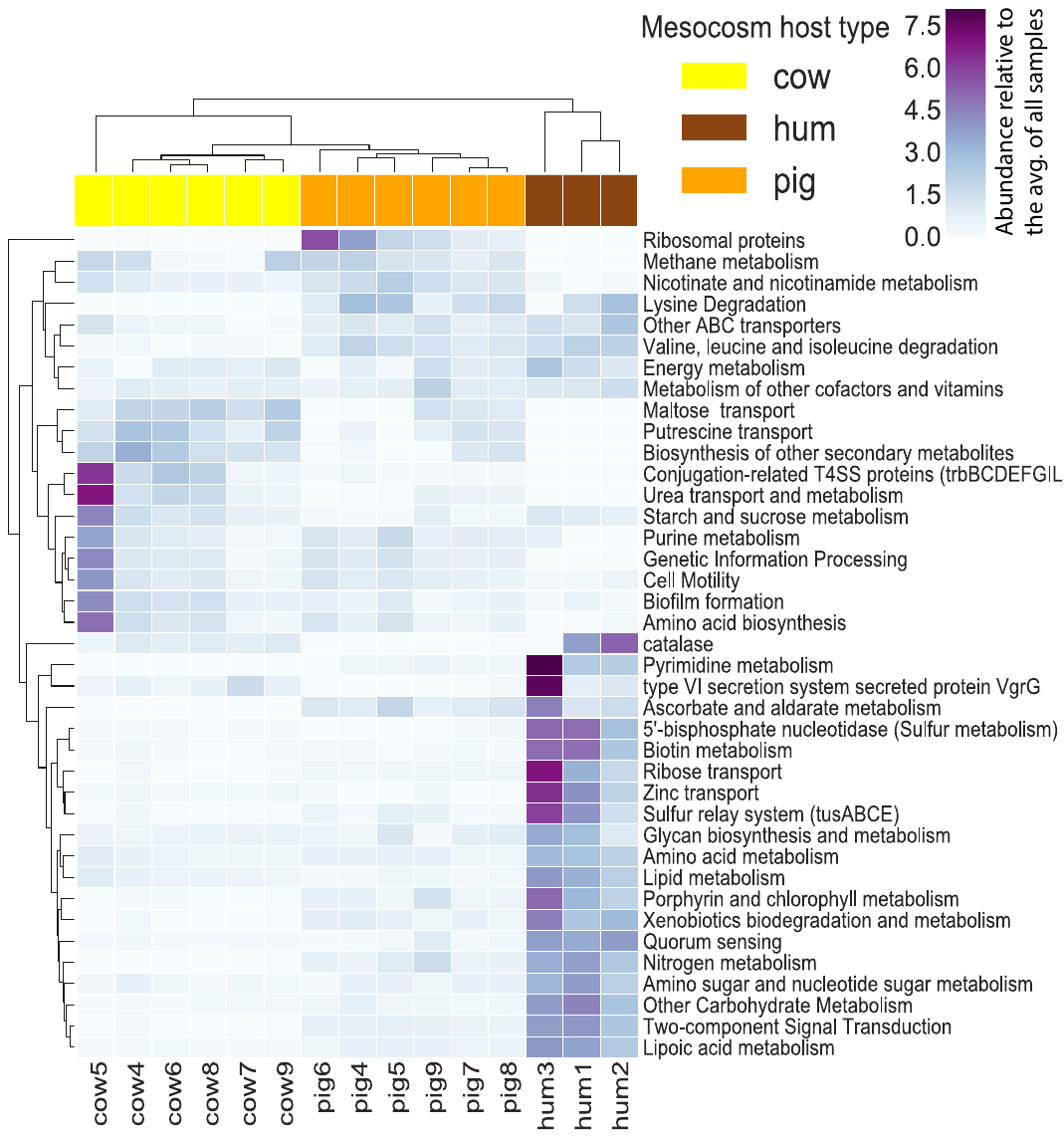

**Figure S13: Gene functions enriched in the cow, pig, or human fecal metagenomes.** The heatmap shows the KEGG functions (rows) that were differentially abundant between the different host fecal metagenomes (columns) with P_adj_ < 0.05 as determined by DESeq2 analysis. Color scale indicates the abundance relative to the average across all metagenome samples.

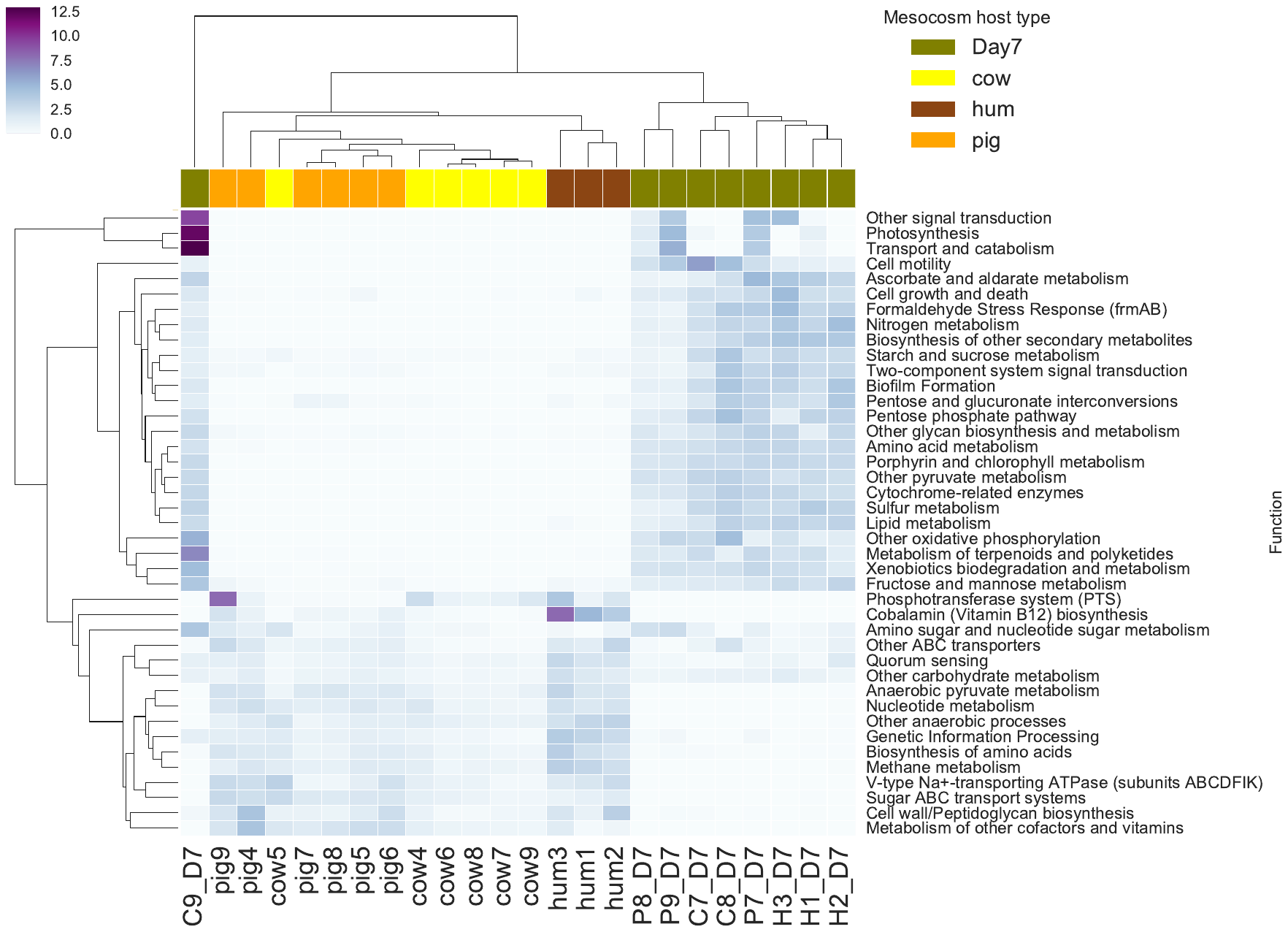

**Figure S14: Gene functions enriched between the host fecal and D7 mesocosm metagenomes.** The heatmap shows the KEGG functions (rows) that were differentially abundant between the different host types (columns) with P_adj_ < 0.05 as determined by DESeq2 analysis. Color scale indicates the abundance relative to the average across all metagenome samples.

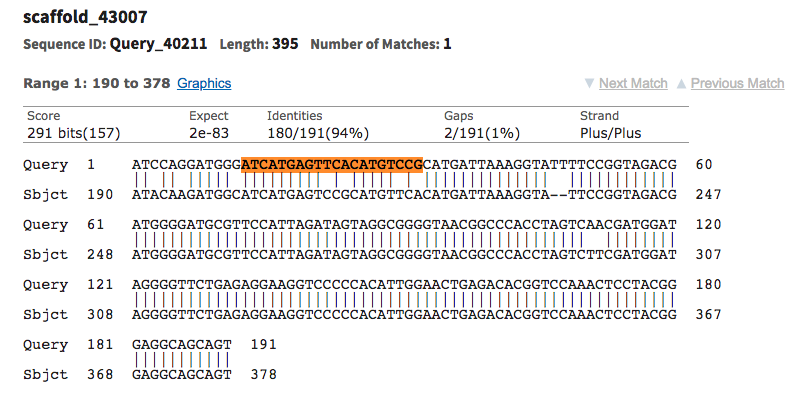

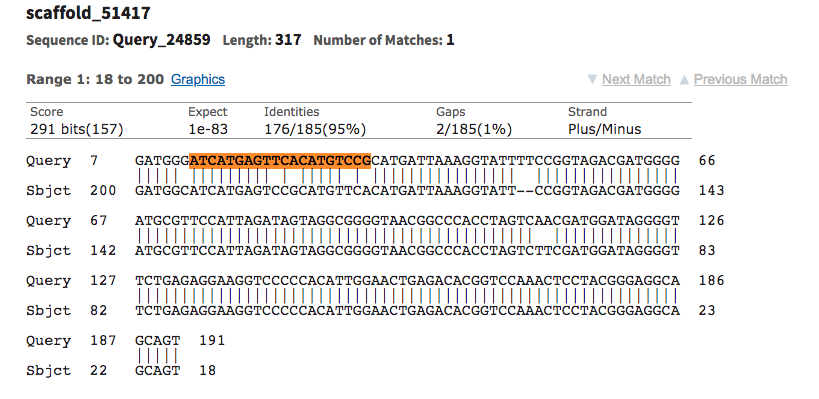

**Figure S15: Alignment of human fecal contigs to reference *B. dorei* 16S rRNA gene sequence.** Contigs from hum1 (top) and hum3 (bottom) fecal assemblies with best match (based on blastn search) to the 16S rRNA gene from the *B. dorei* reference genome (Table 1) and containing perfect matches to the HF183 assay reverse primer and probe (highlighted in yellow) but with mismatches to the forward primer (highlighted in orange).

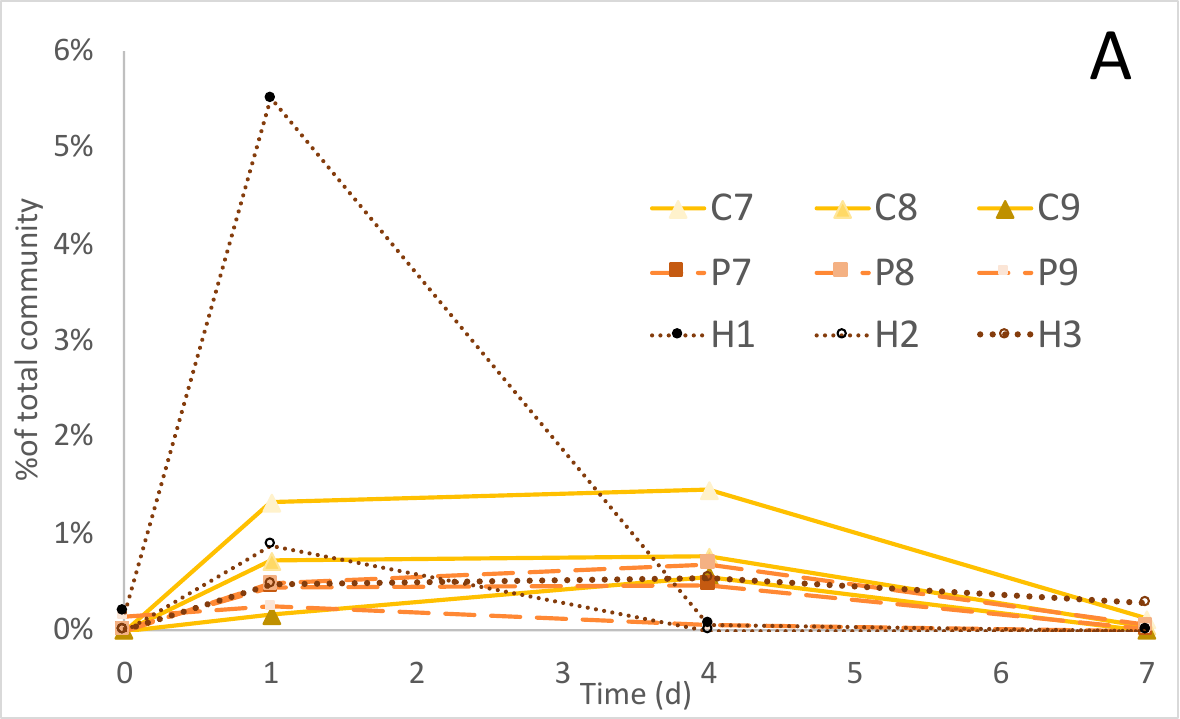

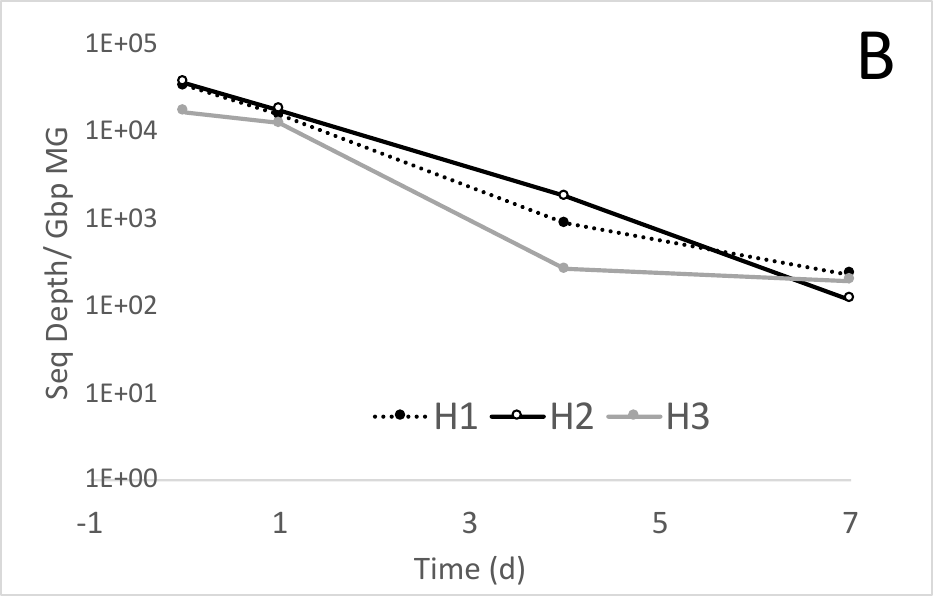

**Figure S16: Decay of *E. coli* and CrAssphage reference genomes in mesocosm metagenomes. (A)** Abundance of the common enteric commensal strain, *Escherichia coli* HS (accession: NC_009800.1), in the fecal mesocosms over time. Abundance is reported as % of total bacterial community (i.e., TAD80 divided by genome sequencing depth; details in the main text). (**B**) Abundance of the human mitochondrial genome (accession: J01415.2) in the human fecal mesocosms expressed as sequencing depth (i.e., TAD80 at >95%ID) divided by metagenomic dataset size in Gbp.

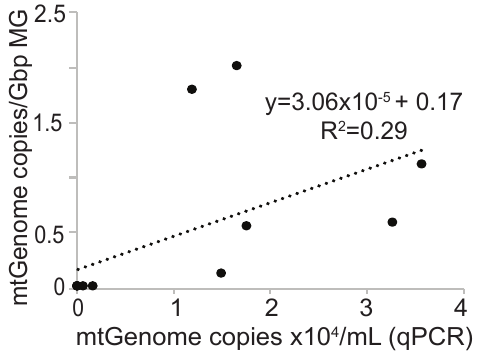

**Figure S17: Correlation between qPCR and metagenome-based abundance estimates of MST markers and their reference genome counterparts.** Human mitochondrial DNA (mtGenome) expressed as the number of gene copies per mL versus the relative abundance of a reference human mtGenome in the metagenome (expressed as TAD80 divided by library size in Gbp).

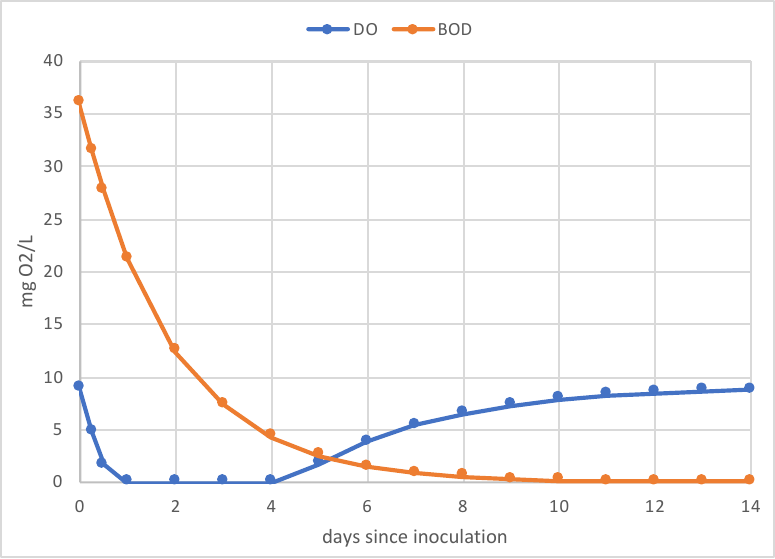

**Figure S18: Modeling dissolved O_2_ (DO) and biological oxygen demand (BOD) in the mesocosms over time.** The Streeter-Phelps equation was used to model BOD depletion and DO re-aeration in the mesocosms assuming total BOD of feces in the tanks is 36 mg/L (36), the water is fully saturated with O_2_ at 20 °C (9 mg/L) and instantaneous O_2_ diffusion into the dialysis bags. Also, a BOD utilization rate of 0.53 day^-1^ and a re-aeration rate of 0.5 day^-1^.

19.	UniProt Consortium, O’Donovan C, Magrane M, Alpi E, Antunes R, Bely B, Bingley M, Bonilla C, Britto R, Bursteinas B, Bye-A-Jee H, Cowley A, Silva AD, Giorgi MD, Dogan T, Fazzini F, Castro LG, Figueira L, Garmiri P, Georghiou G, Gonzalez D, Hatton-Ellis E, Li W, Liu W, Lopez R, Luo J, Lussi Y, MacDougall A, Nightingale A, Palka B, Pichler K, Poggioli D, Pundir S, Pureza L, Qi G, Renaux A, Rosanoff S, Saidi R, Sawford T, Shypitsyna A, Speretta E, Turner E, Tyagi N, Volynkin V, Wardell T, Warner K, Watkins X, Zaru R, Zellner H, Xenarios I, Bougueleret L, Bridge A, Poux S, Redaschi N, Aimo L, Argoud-Puy G, Auchincloss A, Axelsen K, Bansal P, Baratin D, Blatter M-C, Boeckmann B, Bolleman J, Boutet E, Breuza L, Casal-Casas C, Castro E de, Coudert E, Cuche B, Doche M, Dornevil D, Duvaud S, Estreicher A, Famiglietti L, Feuermann M, Gasteiger E, Gehant S, Gerritsen V, Gos A, Gruaz-Gumowski N, Hinz U, Hulo C, Jungo F, Keller G, Lara V, Lemercier P, Lieberherr D, Lombardot T, Martin X, Masson P, Morgat A, Neto T, Nouspikel N, Paesano S, Pedruzzi I, Pilbout S, Pozzato M, Pruess M, Rivoire C, Roechert B, Schneider M, Sigrist C, Sonesson K, Staehli S, Stutz A, Sundaram S, Tognolli M, Verbregue L, Veuthey A-L, Wu CH, Arighi CN, Arminski L, Chen C, Chen Y, Garavelli JS, Huang H, Laiho K, McGarvey P, Natale DA, Ross K, Vinayaka CR, Wang Q, Wang Y, Yeh L-S, Zhang J. 2017. UniProt: the universal protein knowledgebase. Nucleic Acids Res 45:D158–D169.

25.	Morton JT, Marotz C, Washburne A, Silverman J, Zaramela LS, Edlund A, Zengler K, Knight R. 2019. Establishing microbial composition measurement standards with reference frames. Nat Commun 10.
